# The vagus nerve drives cognitive function through the default mode network

**DOI:** 10.64898/2026.10.08.757463

**Authors:** Musashi Yamakawa, Tateki Togashi, Haruki Kawagoe, Reina Shiratori, Fumiya Fukushima, Takuya Urushihata, Ziyan Xie, Tasuku Kayama, Goro Ikeda, Wenfeng Xue, Tuo Xiaoying, Shinpei Kawaoka, Wang Hongbo, Motoko Maekawa, Yuji Owada, Kaoru Seiriki, Hitoshi Hashimoto, Ayumu Konno, Hirokazu Hirai, Yousuke Tsuneoka, Akiko Satoh, Hiroyuki Igarashi, Yusaku Iwasaki, Nariko Arimura, Nahoko Kuga, Takuya Sasaki

## Abstract

The afferent vagus nerve transmits interoceptive signals to the brain but a systematic understanding of how vagal signals affect brain-wide neuronal dynamics remains elusive. Here, comprehensive spatiotemporal analyses across the entire mouse brain revealed that vagus nerve stimulation activated neuronal ensembles within the default mode network (DMN). In contrast, vagal-independent mechanisms predominantly affected the salience network. The vagal-DMN association was mediated by locus coeruleus neuronal subpopulations. Selective activation or silencing of vagal-responsive DMN neuronal ensembles replicated or abolished VNS-induced enhancement of cognitive behaviors and progressive problem-solving behaviors. These findings establish a unifying framework for how the vagus nerve drives cognitive functions by engaging the DMN.

## Introduction

The afferent vagus nerve (VN) ascending from the body to the brain serves as a primary pathway conveying interoception, the sense of the body’s internal state. The VN plays a particularly essential role in regulating emotional valence ^1–5^ and vagus nerve stimulation (VNS) has been shown to improve affective states, exerting anxiolytic and antidepressant effects ^6–8^, as well as to profoundly influence learning, memory, and decision-making ^9–21^. A growing body of evidence suggests that VNS-driven brain functions involve coordinated interplay between widespread cortical networks and neuromodulatory systems, such as locus coeruleus-noradrenergic signaling ^22^ and basal forebrain-cholinergic signaling ^12,17,23^. However, a unified understanding of the neurophysiological mechanisms, spanning molecular, cellular, and whole-brain systems levels, by which the VN modulates higher cognitive function remains elusive.

To address these whole-brain systems-level questions, insights from cognitive neuroscience are instructive, as three major brain networks have been well characterized: the default mode network (DMN) ^24–31^, which operates during rest; the salience network (SN), which activates in response to salient stimuli; and the central executive network, which mediates executive functions. In particular, the DMN plays a crucial role in cultivating memory and intelligence through enhanced self-referential thinking and mind-wandering ^32^, which may underlie systematic mechanisms responsible for VN-related brain functions ^33^. The DMN has also been well characterized in rodent animals with greater inter-subject homogeneity ^34–41^, enabling robust invasive measurements and targeted manipulations.

Building on this cognitive framework, we aimed to systematically and comprehensively unveil the mechanisms and significance of vagal influences on the brain by employing a comprehensive suite of approaches, including whole-brain histological analysis, spatial transcriptomics, functional magnetic resonance imaging (fMRI), and forebrain-wide local field potential (LFP) recordings, in mice. The causal role of higher-order cortical networks at the neuronal level in cognitive function was assessed through targeted manipulation of VNS-recruited neuronal ensembles.

## Results

### Whole-brain analysis of neuronal activation by VNS

We implanted a cuff-shaped electrode on the left cervical VN and applied VNS (0.8 mA, 0.1 ms, 20 Hz, 600 pulses; 30 s on / 30 s off for 3 h) in mice (Fig. 1a) ^3,4,42^. First, we tested whether this VNS protocol was sufficient to affect cognitive behavioral patterns in typical memory and social tests. In a Y-maze alternation test, VNS significantly increased the percentage of correct spontaneous alternations without altering total arm entries at 0–30 min post-stimulation, compared with sham-operated mice with implanted electrodes but no stimulation (Fig. 1b; *n* = 22 and 20 mice; entry: *Z* = 0.82, *P* = 0.41; alternation: *Z* = 2.81, *P* = 0.0050, Mann–Whitney *U* test), suggesting enhanced spatial working memory. In a novel object recognition test, VNS applied before and after the training and before the test phases (24 hours after) significantly increased discrimination index (based on exploration time for the novel and familiar objects) while decreasing total object exploration time (Fig. 1c; *n* = 19 and 20 mice; exploration: *Z* = 2.30, *P* = 0.021; index: *Z* = 2.29, *P* = 0.022, Mann–Whitney *U* test), suggesting enhanced long-term recognition memory. In a social interaction test, VNS significantly increased social interaction ratios defined in an interaction zone, while decreasing time in corner zones (Fig. 1d; *n* = 22 and 19 mice; corner: *Z* = 1.98, *P* = 0.048; ratio: *Z* = 2.29, *P* = 0.022, Mann–Whitney *U* test), suggesting enhanced sociality. These results confirm that our VNS protocol is sufficient to create brain activity states to enhance cognitive behavioral performance.

**Figure 1.**
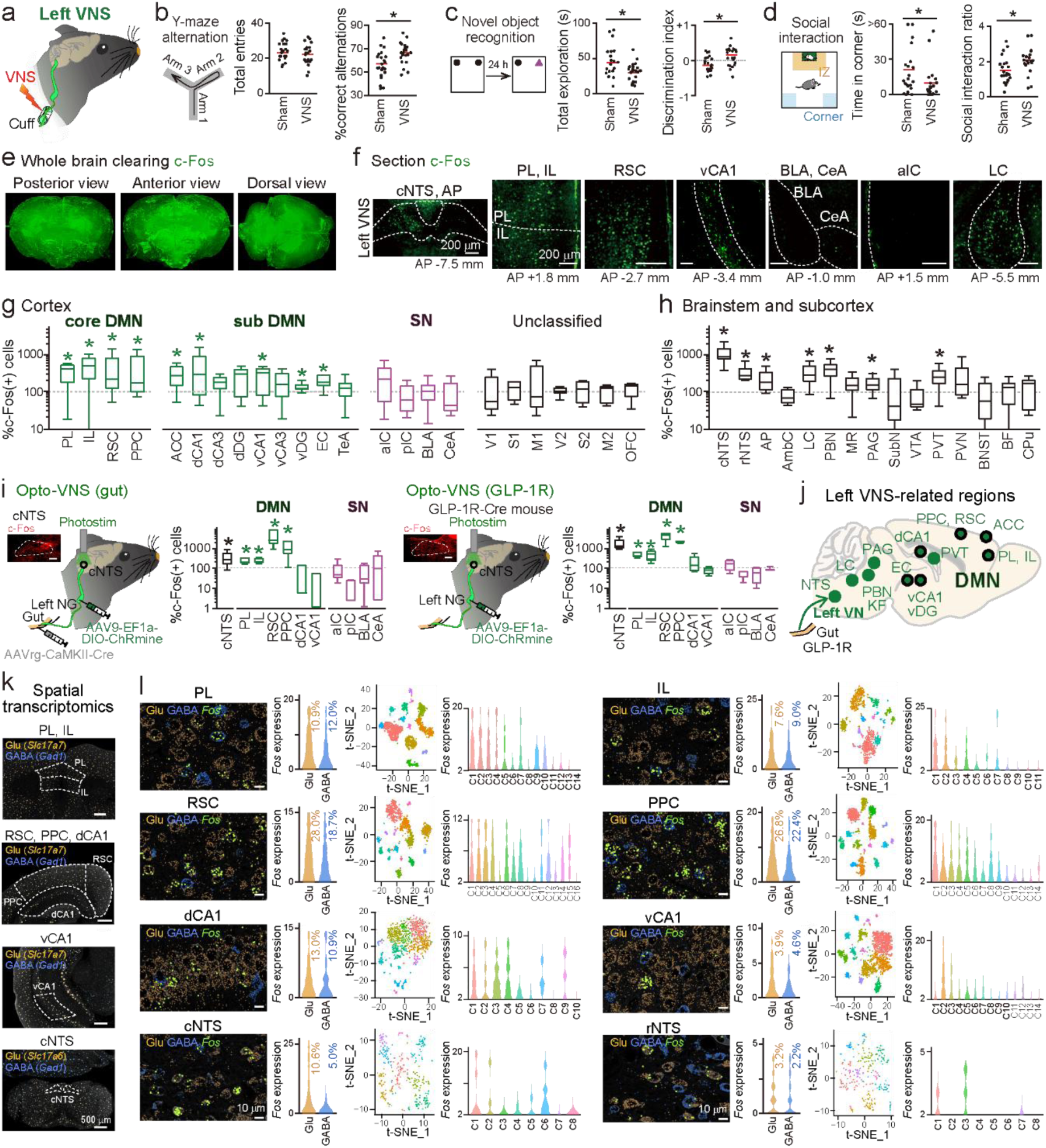
VNS induces neuronal c-Fos expression in the DMN. **a** The left VN was activated by VNS. **b** The total number of arm entries and the percentage of correct spontaneous alternations in a Y-maze test. Each red line represents the average. *n* = 22 and 20 mice, *\*P* = 0.0050, Mann–Whitney *U* test. **c** Total exploration time of two objects (novel: magenta triangle; familiar: black circle) and discrimination index in a novel object recognition test. *n* = 19 and 20 mice, *\*P* = 0.021, \**P* = 0.022, Mann–Whitney *U* test. **d** Total time spent in the corner zone and social interaction ratio in a social interaction test. *n* = 22 and 19 mice, *\*P* = 0.048, \**P* = 0.022, Mann–Whitney *U* test. **e** Whole-brain c-Fos immunostaining after VNS visualized by optical clearing. **f** Coronal sections showing c-Fos expression in representative brain regions following VNS. **g** The percentages of VNS-induced c-Fos-positive neurons in each brain region classified as the DMN and SN, compared with sham control (*n* = 7–9 mice). Box plots show the center line as the median, box limits as the 25th percentile and the 75th percentile, and whiskers as minimum to maximum values that are not outliers. \**P* < 0.05, defined from mean values of 10000 bootstrapped samples. All statistical values are shown in Table 1. **h** Same as g but for the other brainstem and subcortical regions. **i** (Left) Afferent vagal fibers innervating the gut were selectively labeled with ChRmine and activated by photostimulation in the cNTS (*n* = 6 mice). The inset shows c-Fos expression in the cNTS (Scale bar = 200 μm). (Right) GLP-1R-positive afferent vagal fibers were selectively activated (*n* = 4 mice). **j** Schematic illustration of VNS-related brain regions. Black-edged circles represent the DMN. **k** Representative low-magnification images of spatial transcriptomics in each brain region, in which the number of detected marker gene transcripts per cell is indicated by dot size. **l** (From left to right) A magnified image, distribution of the number of *Fos* transcript dots detected per cell in glutamatergic or GABAergic neurons with the percentage of cells in which more than one dot was detected, a t-SNE plot of neuronal clusters, and distribution of *Fos* transcript dots in each cluster (*n* = 2800, 1676, 2349, 1776, 1432, 3072, 692, and 340 cells).

**Table 1.** Statistical results of c-Fos-positive neurons between VNS and sham-operated controls. (From left to right) *P* values calculated by bootstrap method based on data shown in Fig. 1g, 1h (*n* = 7–9 mice), Fig. 1i (*n* = 8 sham and 6 Opto-VNS (gut) mice; *n* = 8 sham and 4 Opto-VNS (GLP-1R) mice), Fig. 3c, 3d (*n* = 6–7 mice sham and 6–10 LPS mice), and Fig. 4l (*n* = 7–9 mice). * indicates a significant increase; # indicates a significant decrease (*P* < 0.05).

| Brain region | *P* values defined by a bootstrapped method |  |  |  |  |
| --- | --- | --- | --- | --- | --- |
|  | Vagal pathway (VNS) | Opto-VNS (gut) | Opto-VNS (GLP-1R) | Vagal-independent pathway (LPS in vagotomy) | Vagal pathway with chemogenetics |
| PL | 0.0092* | 0.047* | 0.0023* | 0.25 | 0.059 |
| IL | 0.0076* | 0.012* | 0.0071* | 0.086 | 0.051 |
| RSC | 0.039* | 0.022* | 0.0028* | 0.96 | 0.085 |
| PPC | 0.023* | 0.048* | 0.00040* | 0.36 | 0.70 |
| ACC | 0.045* | - | - | 0.70 | - |
| dCA1 | 0.048* | 0.94 | 0.14 | 0.52 | 0.10 |
| dCA3 | 0.091 | - | - | 0.85 | - |
| dDG | 0.098 | - | - | 0.47 | - |
| vCA1 | 0.029* | 0.094 | 0.52 | 0.050 | 0.011 (#) |
| vCA3 | 0.094 | - | - | 0.72 | - |
| vDG | 0.031* | - | - | 0.74 | - |
| EC | 0.044* | - | - | 0.95 | - |
| TeA | 0.56 | - | - | 0.94 | - |
| aIC | 0.89 | 0.90 | 0.16 | 0.00030* | 0.14 |
| pIC | 0.67 | 0.040 (#) | 0.30 | 0.0010* | 0.034 (#) |
| BLA | 0.60 | 0.50 | 0.35 | <0.00010* | 0.0098 (#) |
| CeA | 0.95 | 0.31 | 0.32 | <0.00010* | 0.16 |
| V1 | 0.63 | - | - | 0.49 | - |
| S1 | 0.29 | - | - | 0.094 | - |
| M1 | 0.18 | - | - | 0.76 | - |
| V2 | 0.69 | - | - | 0.17 | - |
| S2 | 0.52 | - | - | 0.66 | - |
| M2 | 0.92 | - | - | 0.017(#) | - |
| OFC | 0.45 | - | - | 0.64 | - |
| cNTS | 0.00070* | 0.041* | 0.012* | 0.79 | - |
| rNTS | 0.0019* | - | - | 0.17 | - |
| AP | 0.035* | - | - | 0.41 | - |
| AmbC | 0.99 | - | - | 0.034* | - |
| LC | 0.013* | - | - | 0.49 | - |
| PBN | 0.0049* | - | - | 0.0075(#) | - |
| MR | 0.14 | - | - | 0.17 | - |
| PAG | 0.035* | - | - | 0.79 | - |
| SubN | 0.76 | - | - | 0.42 | - |
| VTA | 0.69 | - | - | 0.27 | - |
| PVN | 0.050 | - | - | 0.00010* | - |
| PVT | 0.014* | - | - | 0.00060* | - |
| BNST | 0.95 | - | - | 0.12 | - |
| BF | 0.56 | - | - | 0.066 | - |
| CPu | 0.28 | - | - | 0.83 | - |

We next examined which brain regions and cell types were activated by VNS through whole-brain, cellular-level analysis of c-Fos expression, an immediate early gene marker of neuronal activation. Whole-brain clearing revealed that c-Fos-expressing neurons were preferentially localized to discrete regions of the brainstem and forebrain (Fig. 1e). Further quantification was performed in each tissue slice (Fig. 1f and Supplementary Fig. 1a). The caudal and rostral nucleus tractus solitarius (cNTS and rNTS), the primary targets of afferent vagal inputs, were confirmed to show significant increases in c-Fos-positive neurons (Fig. 1f and 1h; *n* = 9 mice each; cNTS: *P* = 7.0 × 10^-4^; rNTS: *P* = 0.0019, defined by a bootstrapped method). In addition, among cortical regions analyzed, significant increases in c-Fos-positive neurons were detected in core components of the rodent DMN ^34–41^, including the prelimbic and infralimbic cortex (PL and IL), retrosplenial cortex (RSC), posterior parietal cortex (PPC), and the DMN subsystem, including the anterior cingulate cortex (ACC), dorsal and ventral hippocampal CA1 (dCA1 and vCA1), ventral dentate gyrus (vDG), and entorhinal cortex (EC) (Fig. 1f and 1g; *n* = 7–9 mice; statistics summarized in Table 1, *P* < 0.05 defined by a bootstrapped method). In contrast, no significant changes were observed in brain regions classified as the rodent SN, including the anterior and posterior insular cortex (aIC and pIC), basolateral and central amygdala (BLA and CeA), or other neocortical regions that are not clearly categorized as either the DMN or SN (Table 1, *P* > 0.05). In the brainstem, significant increases in VNS-induced c-Fos expression were also observed in the area postrema (AP), locus coeruleus (LC), parabrachial nucleus (PBN), periaqueductal gray (PAG), and the paraventricular nucleus of the thalamus (PVT) (Fig. 1h and Table 1; *P* < 0.05). Right VNS with the same stimulation parameters induced cNTS activation but no significant increases in c-Fos-expressing neurons in either DMN or SN regions (Supplementary Fig. 1b; *P* > 0.05).

One major source of afferent vagal signals originates from the gut. To stimulate this organ-specific VN pathway, optogenetic VNS was performed by injecting AAVrg-CaMKII-Cre into the duodenum and jejunum and Cre-dependent AAV9-EF1a-DIO-ChRmine-eGFP into the nodose ganglion (NG), followed by green light (532 nm) photostimulation of the cNTS (Fig. 1i, opto-VNS (gut)). This photostimulation similarly resulted in significant increases in c-Fos expression specifically in core DMN regions (Table 1; *P* < 0.05 defined by a bootstrapped method), demonstrating that gut-derived afferent vagal fibers contribute substantially to DMN activation. More specifically, a subset of gut vagal afferent neurons expresses the GLP-1 receptor (GLP-1R) for responding to GLP-1 secreted by intestinal L cells ^43–45^. To selectively activate this vagal subpopulation, optogenetic VNS was performed in GLP-1R-Cre mice by injecting a Cre-dependent AAV9-EF1a-DIO-ChRmine-eGFP into the NG (Fig. 1i, opto-VNS (GLP-1R)). This photostimulation also significantly increased c-Fos expression within the core DMN regions (Table 1; *P* < 0.05). These results indicate that gut-derived GLP-1R-positive vagal afferent fibers contribute to DMN activation (Fig. 1j).

To characterize the cell types exhibiting VNS-induced c-Fos expression in each brain region, we performed spatial transcriptomic analysis across the DMN regions (Fig. 1k). In all regions examined, *Fos* transcripts were detected in subsets of both glutamatergic neurons (defined by *Slc17a7* expression in the cortex and *Slc17a6* expression in the NTS) and GABAergic neurons (defined by *Gad1* expression) (Fig. 1l; *n* = 2800 (PL), 1676 (IL), 2349 (RSC), 1776 (PPC), 1432 (dCA1), 3072 (vCA1), 692 (cNTS), and 340 (rNTS) cells). Transcripts of two other immediate early genes, *Arc* and *Egr1*, were similarly expressed across both excitatory and inhibitory cell types (Supplementary Fig. 2b and Supplementary Data 1; A complete list of gene expression in all analyzed cells is provided in Supplementary Data 2). Furthermore, the number of *Fos* transcripts varied substantially across the 8–16 gene-expression-defined neuron clusters identified by t-distributed stochastic neighbor embedding (t-SNE) analysis (Supplementary Data 1), suggesting that VNS-induced c-Fos expression occurs in specific gene-defined neuronal clusters.

### Neuronal network dynamics modulated by the VN

We further examined VNS-induced whole-brain dynamics by measuring resting-state blood oxygenation level-dependent (BOLD) signals using mouse fMRI ^26,36^ (Fig. 2a and 2b). In both the DMN and SN, VNS did not significantly alter mean amplitude of low-frequency fluctuations (mALFF, 0.01–0.1 Hz), a measure of regional spontaneous BOLD activity ^46,47^ (Fig. 2c and Supplementary Fig. 3a; *n* = 7 mice; DMN: *t*_6_ = 0.78, *P* = 0.47; SN: *t*_6_ = 1.55, *P* = 0.17, paired *t*-test), but it significantly increased functional connectivity specifically within the DMN, but not the SN, as measured by Fisher z-transformed correlation coefficients (z-values) of BOLD temporal correlations (*n* = 7 mice; DMN: *t*_6_ = 3.27, *P* = 0.017; SN: *t*_6_ = 0.35, *P* = 0.73, paired *t*-test). No significant changes in absolute z-values, which reflect the magnitude of connectivity regardless of correlation direction (positive or negative), were observed in either the DMN or SN (*n* = 7 mice; DMN: *t*_6_ = 1.15, *P* = 0.30; SN: *t*_6_ = 0.67, *P* = 0.53, paired *t*-test). These results demonstrate that VNS specifically enhances DMN functional connectivity. We next tested how the deletion of the left VN (vagotomy) affects fMRI signals. Vagotomized mice showed significantly lower absolute z-values of functional connectivity in the DMN and significantly higher absolute z-values in the SN, compared with naïve mice (Fig. 2d; DMN: *t*_23_ = 3.70, *P* = 0.0012; SN: *t*_23_ = 2.93, *P* = 0.0074, Student’s *t*-test), whereas no significant differences in mALFF and z-values of functional connectivity were observed (Fig. 2d and Supplementary Fig. 3b; *n* = 18 and 7 mice; mALFF: DMN: *t*_23_ = 0.60, *P* = 0.56; SN: *t*_23_ = 1.46, *P* = 0.16; z-values: DMN: *t*_23_ = 0.33, *P* = 0.75; SN: *t*_23_ = 0.16, *P* = 0.88, Student’s *t*-test). These results demonstrate that the left VN is required for sustaining certain functional connectivity within the DMN.

**Figure 2.**
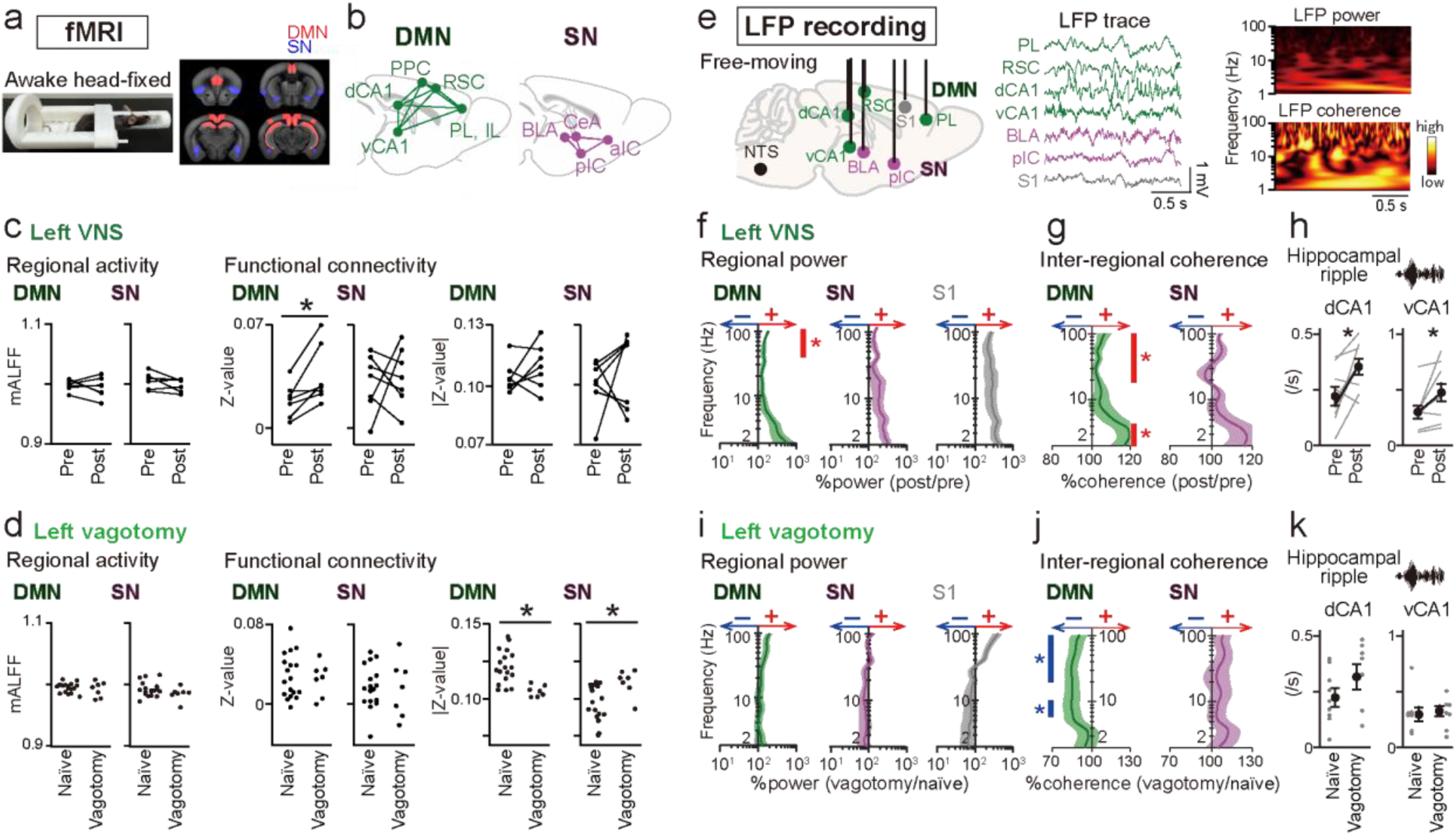
Network activity states induced by the vagal pathway. **a** Resting-state fMRI was performed from awake head-fixed mice. **b** DMN and SN regions analyzed from fMRI data. **c** (Left) Regional activity as mALFF in the DMN or SN regions before and after VNS (*n* = 7 mice). Each line represents each mouse. Individual regions are shown in Supplementary Fig. 3. (Middle and right) Functional connectivity as z-values and absolute z-values. \**P* = 0.017, paired *t*-test. **d** Comparisons of fMRI results from naïve and vagotomized mice (*n* = 18 and 7 mice; DMN: \**P* = 0.0012; SN: \**P* = 0.0074, Student’s *t*-test). Each dot represents each mouse. **e** (Left) LFP signals were recorded from DMN and SN regions. (Right) A PL LFP wavelet spectrum and a PL-RSC coherence spectrum. **f** The percentages of LFP power averaged over the DMN, SN, and S1, after VNS relative to pre-VNS baseline plotted as a function of frequency band (2–100 Hz) (*n* = 9 mice). Shaded areas represent SEM. Red bars indicate frequencies with significant increases (*q* < 0.05, paired *t*-test with FDR correction). **g** The percentages of inter-regional LFP coherence averaged over the DMN and SN after VNS relative to pre-VNS baseline plotted as a function of frequency band (*n* = 9 mice). **h** The event frequencies of dCA1 and vCA1 ripples in pre- and post-VNS periods (dCA1: *n* = 9 mice, \**P* = 0.015; vCA1: *n* = 8 mice, *\*P* = 0.045, paired *t*-test). **i, j** The percentages of LFP power and inter-regional coherence in vagotomized mice relative to naïve mice (*n* = 9 and 7 mice). Blue bars indicate frequencies with significant decreases (*q* < 0.05, Student’s *t*-test with FDR correction). **k** The event frequencies of dCA1 and vCA1 ripples (*n* = 9 and 7 mice).

To further examine VNS-induced neuronal dynamics with higher temporal resolution, we recorded ongoing LFP signals from DMN regions including the PL, RSC, dCA1, and vCA1, and SN regions including the BLA and pIC, and the S1 as a reference area, from resting mice (Fig. 2e). VNS significantly increased gamma (40–100 Hz) LFP power in DMN regions (Fig. 2f and Supplementary Fig. 4b; *n* = 9 mice, *q* < 0.05, paired *t*-test with FDR correction), whereas such significant changes were not observed in the SN regions and S1 (*q* > 0.05). LFP coherence at the delta (2–4 Hz) and gamma (40–100 Hz) bands in the DMN regions was also significantly increased by VNS (Fig. 2g and Supplementary Fig. 4c; *q* < 0.05, paired *t*-test with FDR correction). These VNS-induced changes in LFP coherence patterns are considered to occur in parallel with the increases in functional connectivity observed in BOLD signals. In addition, the event frequencies of both hippocampal dCA1 and vCA1 ripples (150–250 Hz), representing neuronal synchronization related to memory processing ^48^ and DMN activation ^49,50^, were significantly increased by VNS (Fig. 2h; dCA1: *n* = 9 mice, *t*_8_ = 3.09, *P* = 0.015; vCA1: *n* = 8 mice, *t*_7_ = 2.44, *P* = 0.045, paired *t*-test). Taken together, VNS specifically enhanced LFP power, coherence, and hippocampal ripples within DMN regions.

In contrast, vagotomy did not significantly change regional LFP power and the frequencies of hippocampal ripples, compared with naïve mice (Fig. 2i and Supplementary Fig. 4d: *q* > 0.05, Student’s *t*-test with FDR correction; Fig. 2k: dCA1: *n* = 9 and 7 mice, *t*_14_ = 1.87, *P* = 0.083; vCA1: *n* = 8 and 7 mice, *t*_13_ = 0.23, *P* = 0.82, Student’s *t*-test). Contrary to the effects of VNS, inter-regional LFP coherence at the theta (6–10 Hz) and beta-gamma (20–100 Hz) bands in the DMN regions was significantly decreased in vagotomized mice (Fig. 2j and Supplementary Fig. 4e; *q* < 0.05, Student’s *t*-test with FDR correction), whereas such significance was not observed in the SN. These results suggest that the VN is necessary for sustaining LFP theta and gamma coherence within the DMN.

### VN-independent mechanisms activate the SN

In addition to the VN, interoceptive signals are conveyed to the brain via the humoral pathways through the bloodstream, and peripheral sensory nerves. We examined how activation of these VN-independent pathways cooperatively affects whole-brain activity patterns. To collectively activate VN-independent mechanisms, lipopolysaccharide (LPS; 1 mg/kg, i.p.) was administered after vagotomy (Fig. 3a). We found that this condition significantly increased c-Fos-positive neurons in SN regions, but not in any DMN regions (Fig. 3b–3c; *n* = 6–10 LPS mice; *P* < 0.05 defined by a bootstrapped method, Table 1), demonstrating that VN-independent mechanisms are primarily associated with the SN (Fig. 3e), in contrast to the effects of VNS (Fig. 1j). In addition, significant increases in c-Fos expression were also observed in the AmbC, PVN, and PVT (Fig. 3d and Table 1; *P* < 0.05).

**Figure 3.**
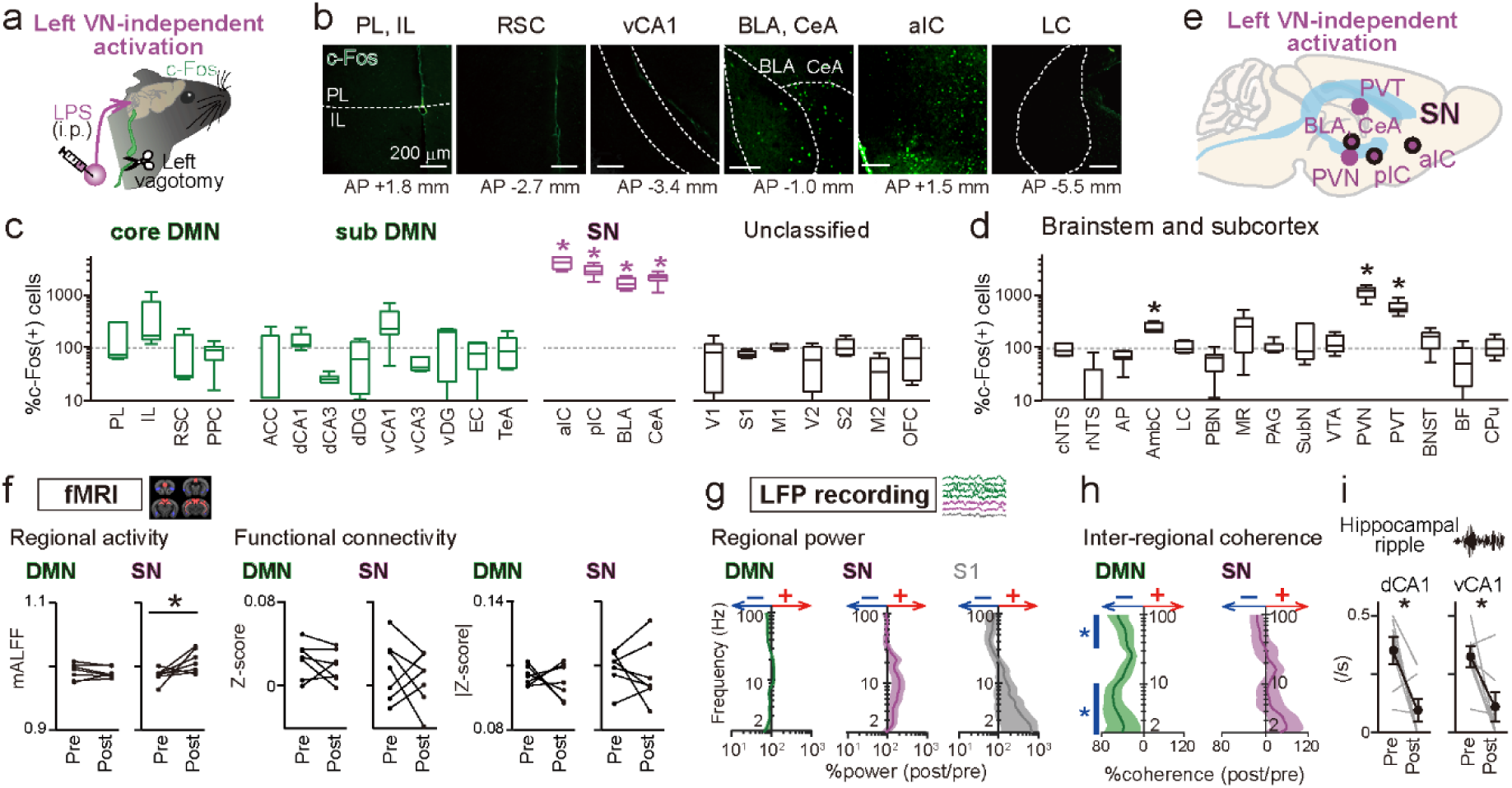
Vagal pathway-independent mechanisms activate the SN. **a** Vagal-independent mechanisms were activated by LPS administration after vagotomy. **b** Images of c-Fos expression in representative brain regions. **c** The percentages of c-Fos-positive neurons in each brain region classified as the DMN and SN, compared with sham control (*n* = 6–10 mice). Box plots show the center line as the median, box limits as the 25th percentile and the 75th percentile, and whiskers as minimum to maximum values that are not outliers. * indicates significant increase (*P* < 0.05), defined from mean values of 10000 bootstrapped samples. All statistical values are shown in Table 1. **d** Same as c but for the brainstem and subcortical regions. **e** Schematic illustration of brain regions activated by LPS after vagotomy. Black-edged circles are SN regions. **f** Changes in fMRI signals in DMN and SN regions by LPS after vagotomy (*n* = 7 mice; \**P* = 0.019, paired *t*-test). **g, h** The percentages of LFP power and inter-regional coherence after LPS relative to pre-LPS baseline plotted as a function of frequency band (2–100 Hz) (*n* = 7 mice). Shaded areas represent SEM. Blue bars indicate frequencies with significant decreases (*q* < 0.05, paired *t*-test with FDR correction). **i** The event frequencies of dCA1 and vCA1 ripples (dCA1: *n* = 7 mice, \**P* = 0.019; vCA1: *n* = 7 mice, *\*P* = 0.027, paired *t*-test).

In the same condition, fMRI analysis detected significant increases in mALFF specifically in the SN, but not in the DMN (Fig. 3f and Supplementary Fig. 3c; *n* = 7 mice; DMN: *t*_6_ = 0.62, *P* = 0.56; SN: *t*_6_ = 2.72, *P* = 0.019, paired *t*-test), whereas functional connectivity in both the DMN and SN was unchanged (z-values: DMN: *t*_6_ = 0.91, *P* = 0.40; SN: *t*_6_ = 0.69, *P* = 0.52; absolute z-values: DMN: *t*_6_ = 0.55, *P* = 0.60; SN: *t*_6_ = 0.52, *P* = 0.62). These fMRI results also confirm that activation of VN-independent mechanisms specifically affects SN regions. In the same condition, LFP recording revealed that coherence across most frequency bands was significantly decreased within the DMN (Fig. 3h and Supplementary Fig. 4g; *n* = 7 mice, *q* < 0.05, 2–100 Hz, except 10–30 Hz, paired *t*-test with FDR correction), but not in the SN (*q* > 0.05 for all regions, paired *t*-test with FDR correction). No significant changes in LFP power across all frequency bands were detected in the DMN and SN regions (Fig. 3g and Supplementary Fig. 4f; *n* = 7 mice; *q* > 0.05 for all regions, paired *t*-test with FDR correction), The frequencies of dCA1 and vCA1 ripples were also significantly decreased (Fig. 3i; dCA1: *n* = 7 mice, *t*_6_ = 3.17, *P* = 0.019; vCA1: *t*_6_ = 2.90, *P* = 0.027, paired *t*-test). Collectively, activation of VN-independent mechanisms reduces electrophysiological functional connections and hippocampal ripples in the DMN, contrasting with the effects of VNS.

### VNS-activated LC noradrenergic neurons mediate the VN-DMN association

To examine detailed pathways underlying the VN-induced DMN activation, trans-synaptic viral tracing was performed by injecting anterograde trans-synaptic AAV1-hSyn-Cre ^51^ into the NG and Cre-dependent AAV9-CAG-DIO-eGFP into the cNTS (Fig. 4a). GFP-positive axonal fibers from cNTS neurons innervated by the VN (VN-cNTS neurons) were observed in the AP, AmbC, PBN, LC, PVN, PVT, and BNST (Fig. 4a; *n* = 6 mice), whereas no apparent GFP-positive VN-cNTS axons were detected in DMN or SN regions. Among these VN-cNTS neuron-innervating regions, we focused on the LC as a relay region since LC neurons were exclusively activated by VNS ^52–54^ (Fig. 1h), but not VN-independent mechanisms (Fig. 3d), and send extensive neuromodulatory projections to the forebrain ^55,56^. Retrograde AAVrg-mCherry tracing from the LC revealed a substantial overlap between mCherry and VNS-induced c-Fos expression in cNTS neurons, confirming that VNS-induced cNTS neurons project to the LC (Fig. 4b). Trans-synaptic labeling revealed that LC neurons receiving cNTS inputs (cNTS-LC neurons) were dopamine β-hydroxylase (*DBH*)-positive and also showed VNS-induced c-Fos expression (Fig. 4c). Moreover, spatial transcriptomic analysis confirmed that VNS-induced *c-fos*-expressing LC neurons were glutamatergic (*Slc17a6*-expressing) or *Th*-expressing neurons, which are predominantly noradrenergic, but not GABAergic (*Gad1*-expressing) types (Fig. 4d and Supplementary Data 1 and 2; *n* = 1247 cells). These results demonstrate that VNS activates noradrenergic LC neuronal subpopulations mediated by cNTS inputs.

**Figure 4.**
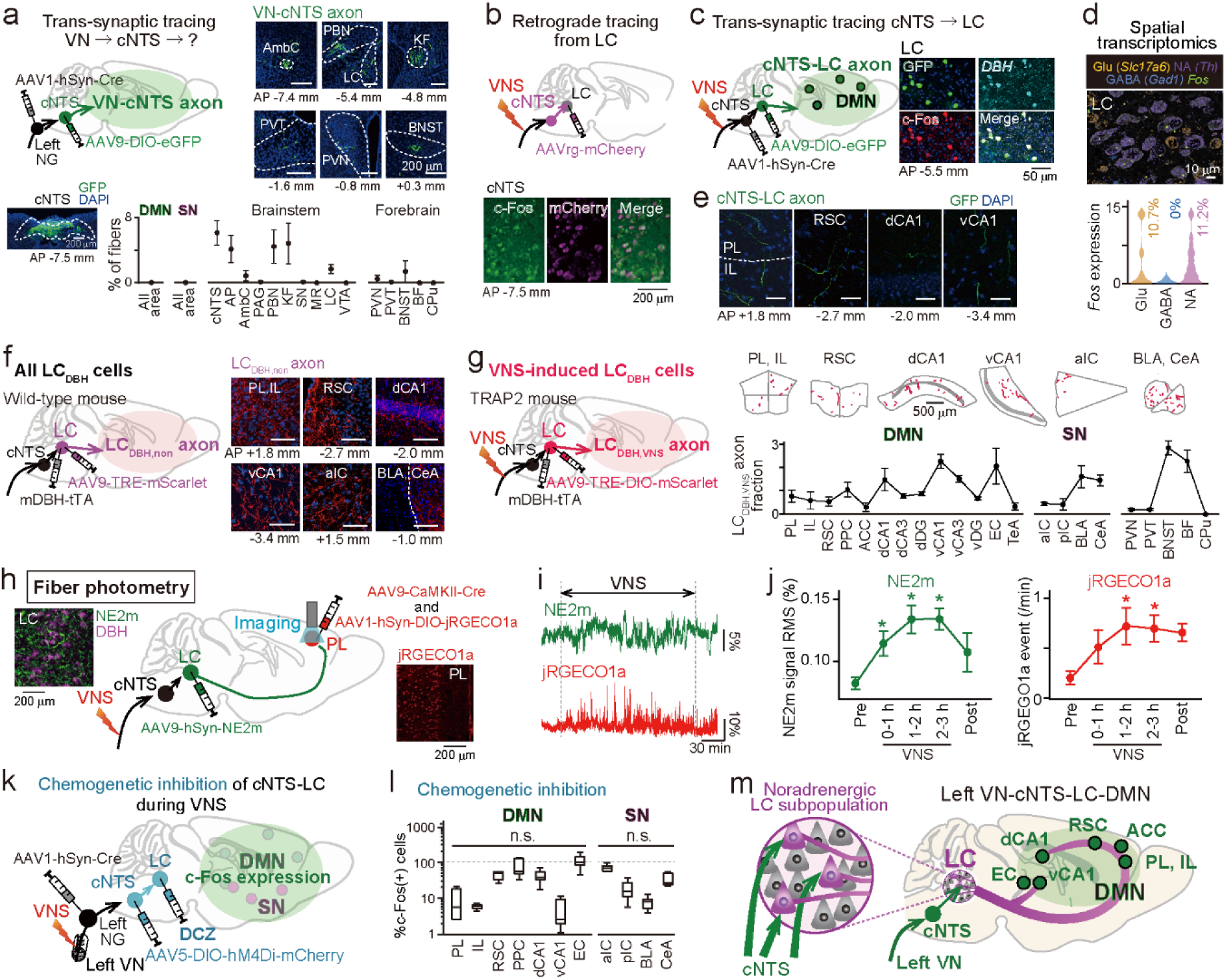
VNS-induced DMN activation is mediated by LC noradrenergic neuronal populations. **a** Axonal projections from cNTS neurons trans-synaptically receiving vagal inputs from the left NG (VN-cNTS; bottom picture) were examined in each brain region. (Top right) Fluorescent images of GFP-expressing VN-cNTS axonal fibers in each brain region. (Bottom) Density of GFP-expressing VN-cNTS axonal projections (*n* = 6 mice). **b** cNTS neurons with axonal projections retrogradely labeled from the LC expressed c-Fos after VNS. **c** LC neurons trans-synaptically labeled from the cNTS (cNTS-LC neurons). These neurons were *DBH*-positive and expressed c-Fos after VNS. **d** Spatial transcriptomics in the LC (*n* = 1247 cells), presented as in Fig. 1l. **e** Axonal projections from cNTS-LC neurons in DMN regions. Scale bars = 200 μm. **f** Axonal projections from all LC DBH-positive neurons (LC_DBH,non_) in each DMN and SN region (*n* = 6 mice). Scale bars = 200 μm. **g** Axonal projections from VNS-induced LC DBH-positive neurons (LC_DBH,VNS_) were detected in each brain region. (Top) Representative detected axons in each region in a mouse. Manually traced axons are shown in red to enhance visibility. (Bottom) LC_DBH,VNS_ axon fractions were computed based on LC_DBH,non_ axon density in each brain region (*n* = 4). **h** LC axonal projections and PL excitatory neurons were labeled with NE2m and jRGECO1a, respectively, and dual fiber photometry imaging was performed from the PL. **i** Simultaneously recorded NE2m and jRGECO1a ΔF/F traces in response to VNS. **j** Changes in the RMS of NE2m signal intensity (left) and the event rates of jRGECO1a signals (right) (*n* = 5 mice). The data are the mean ± SEM. NE2m: Pre versus 0–1 h, \**q* = 0.020; 1–2 h, \**q* = 0.013; 2–3 h, \**q* = 0.0012; jRGECO: Pre versus 1–2 h, \**q* = 0.046; 2–3 h, \**q* = 0.037, paired *t*-test with FDR correction. **k** VN**-**cNTS neurons were labeled with hM4Di, and their axonal projection into the LC was chemogenetically inhibited by local infusion of DCZ. **l** Same as Fig. 1g, but chemogenetic inhibition was performed during VNS (*n* = 9 sham and 7 hM4Di mice), showing no significant increases in any DMN or SN regions (*P* > 0.05). **m** Schematic illustration of VNS-induced activation of DMN regions through LC noradrenergic neuronal subpopulations.

The trans-synaptic tracing from cNTS-LC neurons confirmed substantial axonal projections to DMN regions including the PL, IL, RSC, dCA1, and vCA1 (Fig. 4e). Since the LC is composed of multiple noradrenergic neuronal subpopulations targeting distinct brain regions ^57–61^, we asked whether VNS-responsive noradrenergic neurons in the LC project to DMN regions. First, all LC noradrenergic projections (LC_DBH,non_ axons) were non-selectively labeled by injecting AAV9-mDBH-tTA and AAV9-TRE-mScarlet into the LC, and axonal projection densities were quantified across entire brain regions (Fig. 4f, *n* = 6 mice). This labeling confirmed LC_DBH,non_ axonal projections to an extremely broad range of brain regions, including the DMN and SN, which were subsequently used to quantify the VNS-related axonal projections. Second, VNS-activated LC noradrenergic projections (LC_DBH,VNS_ axons) were selectively labeled by injecting AAV9-mDBH-tTA and AAV9-TRE-DIO-mScarlet into the LC in TRAP2 mice in which 4-hydroxytamoxifen (4-OHT)-inducible Cre recombinase is expressed under the control of the activity-dependent c-Fos promoter ^62,63^ (Fig. 4g; *n* = 4 mice). Axonal projections from LC_DBH,VNS_ neuronal subpopulation in each region were measured by manually tracing axon length. To normalize and evaluate LC_DBH,non_ axon density in each brain region relative to non-selective LC_DBH,non_ axon labeling, the LC_DBH,VNS_ axon fraction in each brain area was quantified as the ratio of (LC_DBH,VNS_ length/total LC_DBH,VNS_ length across all regions) to (LC_DBH,non_ axon density/total LC_DBH,non_ axon density across all regions). Overall, DMN regions showed certain LC_DBH,VNS_ axon fractions, confirming that noradrenergic LC neuron subpopulations activated by VNS substantially projected to DMN regions.

### VNS-activated specific LC neuronal subpopulations mediate the VN-DMN association

Having confirmed that DMN regions receive noradrenergic projections from VNS-activated LC neurons, we next sought to verify its physiological activity patterns by simultaneous fiber photometry recordings of noradrenergic transmission from the LC and neuronal activation in the PL, a typical DMN region. A GRAB_NE2m_ noradrenergic sensor was expressed in LC DBH-positive neurons ^64,65^ by injecting AAV9-hSyn-GRAB_NE2m_ into the LC and a red calcium sensor, jRGECO1a, was expressed in CaMKII-positive neurons in the PL by injecting AAV9-CaMKII-Cre and AAV1-hSyn-DIO-jRGECO1a into the PL (Fig. 4h). An optical imaging fiber was placed in the PL for simultaneous imaging of GRAB_NE2m_ and jRGECO1a signals. Relative to the pre-stimulation baseline, VNS significantly increased the amplitude deflection of GRAB_NE2m_ ΔF/F signals (Fig. 4i and 4j; *n* = 5 mice, Pre versus 0–1 h: *t*_4_ = 4.12, *q* = 0.020; versus 1–2 h: *t*_4_ = 5.19, *q* = 0.013; versus 2–3 h: *t*_4_ = 11.64, *q* = 0.0012; versus Post: *t*_4_ = 1.91, *q* = 0.13, paired *t*-test with FDR correction) and event rates of jRGECO1a ΔF/F signals (Fig. 4j; *n* = 5 mice, Pre versus 0–1 h: *t*_4_ = 2.25, *q* = 0.088; versus 1–2 h: *t*_4_ = 3.16, *q* = 0.046; versus 2–3 h: *t*_4_ = 3.85, *q* = 0.037; versus Post: *t*_4_ = 4.19, *q* = 0.055, paired *t*-test with FDR correction). These results demonstrate that LC noradrenaline release synchronizes with PL activation after VNS, suggesting that VNS-induced PL neuronal activation is mediated by LC noradrenergic projections.

To directly test the contribution of the VN-cNTS-LC pathway to the VNS-induced DMN activation, we performed chemogenetic inhibition of VN-cNTS axonal terminals in the LC by injecting AAV1-hSyn-Cre into the NG and AAV5-DIO-hM4Di-mCherry into the cNTS, followed by local DCZ administration into the LC (Fig. 4k). The chemogenetic inhibition prevented VNS from significantly increasing the proportions of c-Fos-positive neurons in all brain regions classified as either the DMN or SN, compared to sham-operated controls without VNS (Fig. 4l; *n* = 7–9 mice; *P* < 0.05 defined by a bootstrapped method, statistics summarized in Table 1). Taken together, these results demonstrate that VNS-induced neuronal activation in the DMN regions is mediated by specific VN-cNTS-LC noradrenergic neuronal subpopulations (Fig. 4m).

### VNS-induced DMN neurons are necessary and sufficient for enhancing cognitive behavior

We examined whether VNS-induced DMN neurons causally contribute to cognitive functions. In TRAP2 mice, Cre-dependent AAV8-hSyn-DIO-hM4Di-mCherry was simultaneously injected into four DMN regions, including the PL, RSC, dCA1, and vCA1 (Fig. 5a), and a cuff electrode was implanted on the left VN. After two weeks, mice received 4-OHT injection followed by VNS to label VNS-activated neurons with hM4Di (Fig. 5b). After seven days, mice were intraperitoneally administered 1 mg/kg DCZ to inhibit hM4Di-labeled neurons (VNS-DMN inhibition group). VNS was then applied before each behavioral test, using the same protocols as in Fig. 1b–d. In contrast to the enhancing effects observed of VNS (Fig. 1b–d), the VNS-DMN inhibition mouse group showed no significant VNS-induced improvement in cognitive-related behaviors in the Y-maze alternation test (Fig. 5c; *n* = 22 and 6 mice; entry: *Z* = 1.29, *P* = 0.20; alternation: *Z* = 0.61, *P* = 0.54), novel object recognition test (Fig. 5d; *n* = 19 and 6 mice; exploration: *Z* = 2.77, *P* = 0.0056; index: *Z* = 1.37, *P* = 0.17), or social interaction test (Fig. 5e; *n* = 22 and 6 mice; corner: *Z* = 1.60, *P* = 0.11; ratio: *Z* = 0.14, *P* = 0.89, Mann–Whitney *U* test), compared with sham-operated mice without VNS. These results demonstrate that VNS-induced DMN neurons are indeed necessary for VNS to exert its cognitive-enhancing effects.

**Figure 5.**
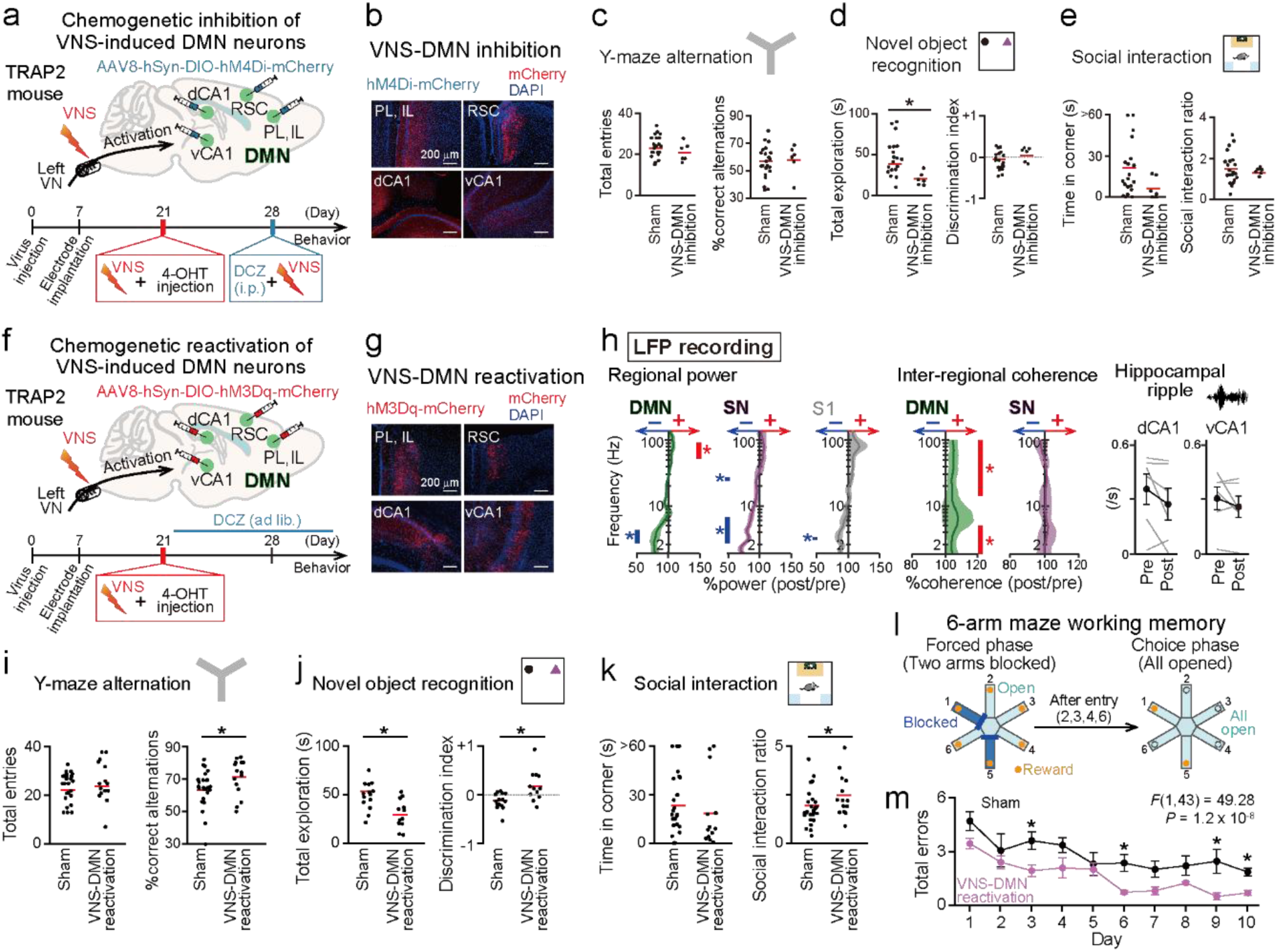
VNS-induced DMN neuronal ensembles enhance cognitive behaviors. **a** Using the TRAP2 system, VNS-induced neuronal ensembles in four DMN regions were labeled with hM4Di (day 21). On a behavioral test day (day 24), DCZ was first applied, and VNS was then applied 30 min after DCZ administration, followed by behavioral testing (VNS-DMN inhibition). The procedure from VNS to the subsequent behavioral testing was the same as in Fig. 1b–d. **b** Pictures of mCherry-expressing neurons in four DMN regions for VNS-DMN inhibition. **c** Y-maze alternation test performance. Each red line represents the average. *n* = 22 and 6 mice. **d** Novel object recognition test performance. *n* = 19 and 6 mice, *\*P* = 0.0056, Mann–Whitney *U* test. **e** Social interaction test performance. *n* = 22 and 6 mice. **f,g** Using the same TRAP2 system, VNS-induced neuronal ensembles were labeled with hM3Dq (VNS-DMN reactivation). **h** Changes in LFP power signals, inter-regional LFP coherence plotted as a function of frequency band, and the frequencies of dCA1 and vCA1 ripples by VNS-DMN reactivation (*n* = 6 mice). Shaded areas represent SEM. Red and blue bars indicate frequencies with significant increases and decreases, respectively (*q* < 0.05, paired *t*-test with FDR correction). **i** Y-maze alternation test performance. *n* = 25 and 15 mice, *\*P* = 0.039, Mann–Whitney *U* test. **j** Novel object recognition test performance. *n* = 15 and 12 mice, *\*P* = 0.0032, *\*P* = 0.0040, Mann–Whitney *U* test. **k** Social interaction test performance. *n* = 27 and 14 mice, *\*P* = 0.029, Mann–Whitney *U* test. **l** Schematic illustration of the six-arm radial maze task. **m** Average number of errors per trial across days. *n* = 4 and 5 mice. \**P* < 0.05, Welch’s *t*-test with Benjamini-Hochberg FDR correction.

Furthermore, we examined whether direct activation of VNS-induced DMN neurons, in the absence of VNS, is sufficient to recapitulate the cognitive-enhancing effects of VNS. With the same protocol as in Fig. 5a, we labeled VNS-activated neurons with hM3Dq by injecting AAV8-hSyn-DIO-hM3Dq-mCherry in TRAP2 mice (Fig. 5f and 5g). During the resting state, LFP recording demonstrated that DCZ administration to these mice (VNS-DMN reactivation) significantly increased gamma (50–100 Hz) LFP power and coherence across a wide frequency range, including the delta and gamma bands, in DMN regions (Fig. 5h and Supplementary Fig. 4h and 4i; *n* = 6 mice, *q* < 0.05, paired *t*-test with FDR correction), whereas such significant changes were not observed in the SN regions (*q* > 0.05). These changes in LFP patterns closely resembled those observed with VNS (Fig. 2f and 2g), validating that VNS-DMN reactivation partially reproduces the VNS-induced neuronal ensemble activity within the DMN. No significant differences in hippocampal ripples were observed following VNS-DMN reactivation (dCA1: *n* = 6 mice, *t*_5_ = 1.66, *P* = 0.16; vCA1: *n* = 6 mice, *t*_5_ = 1.23, *P* = 0.27, paired *t*-test).

Next, VNS-DMN reactivation was constitutively induced by administration of DCZ-containing drinking water. These mice showed significant improvements across all the three tests, compared with sham mice with no reactivation (TRAP2 mice injected with AAV8-hSyn-DIO-hM3Dq-mCherry without DCZ): (1) increased percentage of correct spontaneous alternations in the Y-maze test (Fig. 5i; *n* = 25 and 15 mice; entry: *Z* = 0.59, *P* = 0.56; alternation: *Z* = 2.07, *P* = 0.039), (2) increased discrimination index in the novel object recognition test (Fig. 5j; *n* = 15 and 12 mice; exploration: *Z* = 2.95, *P* = 0.0032; index: *Z* = 2.88, *P* = 0.0040), and (3) increased social interaction ratio in the social interaction test (Fig. 5k; *n* = 27 and 14 mice; corner: *Z* = 1.20, *P* = 0.23; ratio: *Z* = 2.19, *P* = 0.029, Mann–Whitney *U* test). These effects could also simply be attributable to an overall increase in activated neurons within DMN regions. To address this possibility, we tested a random activation group, in which TRAP2 mice injected with Cre-independent AAV8-hSyn-hM3Dq-mCherry (2.0 × 10¹¹ vg/ml) received DCZ to randomly activate a similar proportion of neurons (10–20%) without VNS-induced specificity (Supplementary Fig. 5a and 5b). The random activation mice did not show any significant improvements in the three tests (Supplementary Fig. 5c; Y-maze alternation: *n* = 25 and 6 mice; entry: *Z* = 2.26, *P* = 0.024; alternation: *Z* = 1.03, *P* = 0.30; novel object recognition: *n* = 15 and 5 mice; exploration: *Z* = 1.75, *P* = 0.081; index: *Z* = 0.79, *P* = 0.43; social interaction: *n* = 27 and 5 mice; corner: *Z* = 0.42, *P* = 0.68; ratio: *Z* = 0.18, *P* = 0.86, Mann–Whitney *U* test), compared with sham mice with no reactivation. These results demonstrate that reactivation of VNS-induced neuronal ensembles, but not randomly selected neuronal ensembles, within the DMN specifically trigger cognitive-enhancing behavioral effects, similar to VNS.

We further assessed the effects of DMN reactivation on more complex problem-solving behaviors using a six-arm radial maze task (Fig. 5l). In each trial, four of the six arms were randomly selected and left open, while the remaining two were blocked by barriers. Chocolate milk (50 μl) was placed at the end of all six arms as a reward. Each trial consisted of two phases: in the forced phase, mice were first placed at the central platform and required to visit all four open arms to collect rewards; subsequently, in the choice phase, the barriers were removed, and all six arms became open. A trial was completed when the mouse had visited all six arms. Re-entries into previously visited arms were scored as errors, with the optimal strategy being to visit each arm exactly once. Five trials were conducted per day. In the early days, mice committed frequent errors but progressively reduced error rates across days, reflecting the acquisition of an efficient behavioral strategy and/or improved spatial working memory. Over 10 days of testing, both sham mice with no reactivation and mice with VNS-DMN reactivation showed a significant reduction in errors across days (Fig. 5m; *n* = 4 and 5 mice; *F*(9,171) = 3.46, *P* = 6.0 × 10^-4^; *F*(9,216) = 12.33, *P* = 1.0 × 10^-15^, repeated-measures ANOVA), reflecting cumulative learning over successive days. However, VNS-DMN reactivation mice showed significantly fewer errors than sham mice on days 3, 6, 9, and 10 (*P* < 0.05, Welch’s *t*-test with Benjamini-Hochberg FDR correction), and overall across the entire 10-day period (*F*(1,43) = 49.28, *P* = 1.2 × 10^-8^, repeated-measures ANOVA). These results demonstrate that reactivation of VNS-induced DMN neurons is also sufficient to improve progressive and demanding cognitive behaviors.

## Discussion

Our comprehensive whole-brain analyses demonstrated that left VNS increased c-Fos-positive neurons, inter-regional functional connectivity, gamma power and coherence specifically in the DMN. The vagal-DMN association is supported by noradrenergic projections from specific VNS-responsive neuronal subpopulations in the LC. Selective chemogenetic inhibition of VNS-induced neuronal ensembles in the DMN abolished the cognitive-enhancing effects of VNS, while their reactivation was sufficient to improve cognitive functions. These findings highlight the vagal-DMN axis as a key substrate for enhancing higher-order brain functions.

While human fMRI studies indicate that transcutaneous auricular VNS (taVNS) recruits brain regions involved in the DMN, including the precuneus, medial prefrontal cortex, posterior cingulate cortex, and parahippocampal regions ^53,66–68^, other studies have reported pronounced effects extending beyond the DMN to SN regions, suggesting considerable inconsistency across studies ^67–69^. Our studies in rodents, benefiting from direct invasive measurements, controlled experimental conditions and individual homogeneity, provide robust evidence that VNS specifically engages the DMN at the cellular level and enhances cognitive and intellectual behavioral performance. These results are in accordance with the established role of the DMN in supporting higher-order cognitive processes, including abstract reasoning, problem-solving, and internally directed thought. We also demonstrated that VNS increased hippocampal ripples, which are shown to be linked to DMN generation ^49,50^ and serve as neurophysiological substrates coordinating widespread cortical activity across brain networks to support memory processing, mental simulation, and prospective planning ^48,70^. Taken together, these coordinated effects indicate that left VNS promotes an introspective brain state conducive to memory processing and higher-order cognition by engaging the DMN. In contrast, we demonstrated that activation of VN-independent pathways specifically affects the SN while reducing DMN activity. These findings suggest that distinct interoceptive mechanisms are selectively linked to specific cognitive networks (i.e., VN to the DMN, VN-independent to the SN), potentially inducing distinct cognitive modes.

The afferent VN comprises heterogeneous organ-, projection-, and gene-specific subtypes ^71–75^, potentially broadening the range of interoceptive signals conveyed to the brain. Among these VN subtypes, we demonstrated that activation of the gut-derived, GLP-1-positive VN is sufficient to replicate VNS-induced increases in c-Fos-positive neurons in the DMN. Given that GLP-1 signaling is elevated postprandially, reflecting nutrient availability ^43–45,76^, such physiologically replete brain-body states may naturally engage vagal-DMN association. More broadly, various subtypes of the afferent VN likely respond to distinct physiological conditions and may similarly reinforce vagal-DMN coupling.

VNS has been well established to activate the LC in both rodents ^22,52,54^ and humans ^53^, thereby exerting widespread cortical influences ^22^. In the LC, neuronal populations are highly heterogeneous, with individual noradrenergic neurons projecting to distinct brain regions ^57–60^. Here, we identified specific LC neuronal subpopulations responsive to VNS that mediate DMN neuronal activation via noradrenergic projections. These findings reveal a previously unrecognized role for LC-derived noradrenaline. Although the LC noradrenergic system is generally associated with arousal and attention ^22^, our results suggest that VNS-responsive LC-DMN projections rather support introspective memory.

Multiple mechanisms have been proposed to underlie DMN formation, including serotonergic ^77^ and cholinergic ^78^ modulation, dynamic balanced interactions with the SN ^37,38^, and intracortical connectivity within the DMN ^35^. Our analysis did not detect VNS-induced pronounced increases in median raphe-serotonergic or basal forebrain-cholinergic neuronal activity. We consider that the contribution of cholinergic signaling may be more prominent under temporally constrained conditions dependent on external sensory stimuli or task demands ^12,17,23^, while serotonergic contributions may be more apparent under chronic conditions ^79^ or pathological conditions such as depression. Our results suggest that, under temporally unconstrained conditions with minimal restriction on spontaneous behavior, specific LC subpopulations mediate the noradrenergic drive through which VNS induces superior cognitive states.

In conclusion, our study identifies the VN as a powerful gateway to superior cognitive functions. From a clinical perspective, accumulating evidence demonstrates that DMN dysfunction contributes to various neurodegenerative and psychiatric disorders, including Alzheimer’s disease, schizophrenia, and depression ^30,77,80^. Targeting the vagal-DMN interoceptive association may therefore provide novel therapeutic strategies rooted in brain-body communication.

## Methods

### Approvals

All experiments were performed with the approval of the animal experimental ethics committee at Tohoku University (approval number: 2022 PhA-004) and in accordance with the NIH guidelines for the care and use of animals.

### Subjects

C57BL/6J mice (eight to ten weeks old) with weights of 29–35 g were used. All wild-type mice were purchased from SLC (Shizuoka, Japan). TRAP2 mice purchased from Jackson Laboratory (*Fos^tm2.1(icre/ERT2)Luo^*/J; stock #030323) were used at eight to ten weeks of age with body weights ranging from 29–35 g. In this study, we used both male and female mice; however, behavioral analyses were restricted to male mice, as behavioral patterns were not fully reproducible in female mice. They were maintained with free access to water and food under a 12-h light/12-h dark schedule under housing conditions at 23 ± 1°C with relative humidity of 50 ± 5% with lights off at 8:00 PM.

### Surgery for vagus nerve stimulation, electrophysiological recordings, and drug and virus injection

Mice were anesthetized with 3% isoflurane gas and then maintained with 1–3% isoflurane gas while lying on their backs. Veterinary ointment was placed on the mouse’s eyes to prevent dryness. For all steps of an incision, the skin was sterilized with betadine and 70% ethanol.

For vagotomy and VNS, an incision was made in the left neck area from the larynx to the sternum, and the bundle including the VN and the carotid artery was isolated. The left cervical VN was isolated from the surrounding tissue and the left carotid artery. Here, we primarily targeted the left VN because (1) the left VN has fewer connections to the sinoatrial node, compared with the right VN, and the left VNS thus induces fewer detrimental side effects on cardiac activity, and (2) the left VNS has been shown to induce anxiolytic effects and affect emotion-related functions ^4^. For vagotomy, the cervical VN bundle was exposed and dissected by surgical micro scissors. For VNS, the isolated cervical VN was enclosed by a custom-made cuff-shaped VN electrode with the electrode contact sites attached to the inside walls of the cylindrical tube (inner diameter = 0.5 mm, electrode area = 0.15 mm^2^, cathode–anode interval = 0.5 mm, and total length 3.0 mm) ^4^. Right VNS was performed using the same protocol.

For LFP recordings, after implantation of the VNS electrode, a 2-cm midline incision was made from the area between the eyes to the cerebellum. Craniotomies with a diameter of ∼1 mm were created above target brain regions using a high-speed drill, and electrodes were implanted into the brain. An electrode assembly was composed of an electrical interface board (EIB-36-PTB, Neuralynx, Inc., Bozeman, MT), an outer cover, and a core body created by 3D printers (UP Plus2) ^3,4,81^ with electrodes (17-μm-wide polyimide-coated platinum-iridium (90/10%) wires (California Fine Wire) plated with platinum to lower the electrode impedances to 200–250 kΩ) for 10 LFP channels and 2 ground/reference (g/r) channels. The electrode assembly was stereotaxically implanted in the left hemisphere at the following coordinates (mm from bregma): PL (AP: 2.0, ML: 0.2, DV: –2.0), RSC (AP: –2.0, ML: 0.2, DV: –0.5), dCA1 (AP: –1.8, ML: 1.4, DV: –1.5), vCA1 (AP: –3.0, ML: 3.5, DV: – 4.6), BLA (AP: –1.0, ML: 3.7, DV: –4.4), pIC (AP: –0.5, ML: 4.5, DV: –4.0), and S1 (AP: – 1.0, ML: 3.8, DV: –2.0). Ground electrodes were located on the cerebellum. The recording device was secured to the skull using stainless steel screws and dental cement. After all surgical procedures were completed, anesthesia was discontinued, and the mice were allowed to awaken spontaneously. Following surgery, each mouse was housed in a transparent Plexiglas cage with free access to water and food. While all mice were implanted with electrodes for simultaneous electrophysiological recordings, some electrodes were implanted outside the target regions or accidentally disrupted during chronic implantations. In these cases, we recorded as many signals as possible simultaneously from the individual mice.

For fMRI, after implantation of the VNS electrode or vagotomy, a custom-made head bar was affixed to the skull using dental cement. The head bar was subsequently used to secure the mice for fMRI experiments.

For optogenetic experiments to test the gut-specific vagal pathway, AAVrg-CaMKII-Cre (the UNC Vector Core service) dissolved in phosphate-buffered saline (PBS) was injected into the duodenum and jejunum in wild-type mice. In addition, a glass pipette (ϕ = 10 µm) was inserted into the left NG and a mixture of Cre-dependent AAV9-EF1a-DIO-ChRmine-eGFP (the UNC Vector Core service) and 0.1% Fast Green (F7252-5G, Sigma-Aldrich, US) dissolved in PBS (total volume ∼2 μl) was injected. Successful injection was confirmed by uniform Fast Green staining throughout the NG. In addition, an optical fiber (ϕ = 200 µm) was implanted into the cNTS. Photostimulation with green light pulses (532 nm, 15.9 mW/mm^2^) was applied. For optogenetic stimulation of GLP-1R-positive vagal afferents, the same manipulations without gut injection were applied to GLP-1R-Cre mice (*Glp1r^tm1.1(cre)Lbrl^/RcngJ*; stock #029283).

For axonal tracing from the NG, the NG was surgically exposed. A glass pipette (ϕ = 50 µm) beveled to 30° using a micropipette grinder (EG402; Narishige, Tokyo, Japan) was inserted into the NG, and a mixture of AAV5-hSyn-mCherry (the UNC Vector Core service) in PBS (total volume ∼2 μl) was injected at an air pressure of 200 hPa.

For trans-synaptic axonal tracing, AAV1-hSyn-Cre (the UNC Vector Core service) was injected into the NG as described above or injected into the cNTS (at the coordinate (mm from obex): AP: –0.5, ML: ±0.2, DV: –0.5) at a rate of 100 nl/min for 1 min using a syringe pump (Muromachi Kikai, Tokyo, Japan). In addition, a glass pipette (ϕ = 10 µm) was inserted into the cNTS or the LC (at an angle of 15°; at the coordinate (mm from bregma): AP: –6.0, ML: ±0.95, DV: –3.1) and Cre-dependent AAV9-CAG-DIO-eGFP (the UNC Vector Core service) was injected as described above. After all injection into the brain tissue, the injection pipette was left in place for 10 min.

For axonal tracing from non-selective LC noradrenergic neuron types (LC_DBH,non_ neurons), AAV9-mDBH-tTA-SV40pA and AAV9-TRE-Gap43pal-mScarlet-WPRE-hGHpA ^57^ were injected into the LC in wild-type mice.

For axonal tracing from VNS-induced LC noradrenergic neurons (LC_DBH,VNS_ neurons), AAV9-mDBH-tTA-SV40pA and AAV9-TRE-DIO-Gap43pal-mScarlet-WPRE-hGHpA ^57^ were injected into the LC in TRAP2 mice. 4-hydroxytamoxifen (4-OHT) dissolved in a 1:9 (v/v) mixture of ethanol and sunflower oil at a concentration of 10 mg/mL was administered as a single intraperitoneal injection at 100 mg/kg, 1 hour prior to VNS. VNS was applied to label TRAPped neurons.

For fiber photometry experiments, AAV9-hSyn-GRAB_NE2m_-WPRE-hGH-polyA (BrainVTA, China) was injected into the LC and a 1:1 mixture of AAV9-CaMKII-GFP-P2A-Cre and AAV1-hSyn-Flex-NES-jRGECO1a-WPRE-SV40, a gift from Douglas Kim & GENIE Project (addgene, 100853-AAV1), was injected into the PL (at the coordinate (mm from bregma): AP: +2.0, ML: 0.0, DV: –1.8).

For chemogenetic experiments to test the NG-cNTS pathway, AAV1-hSyn-Cre was injected into the NG and AAV5-DIO-hM4Di-mCherry (the UNC Vector Core service) was injected into the cNTS as described above. In addition, guide tubes (inner diameter = 0.34 mm and outer diameter = 0.5 mm) were bilaterally implanted into the LC at an angle of 15° (at the coordinate (mm from bregma): AP: –6.0, ML: ±0.95, DV: –3.1). To prevent drying in the guide tubes, a dummy plastic cannula with a diameter of 0.33 mm was inserted into the guide tube until the DCZ injection described below was started.

For chemogenetic experiments to test the VNS-related DMN neuronal ensembles, AAV8-hSyn-DIO-hM4Di-mCherry, AAV8-hSyn-DIO-hM3Dq-mCherry, AAV8-hSyn-hM3Dq-mCherry, or AAV8-hSyn-mCherry (the UNC Vector Core service) was injected into the left hemisphere at the following coordinates (mm from bregma): PL (AP: 2.0, ML: 0.2, DV: –1.5), RSC (AP: –2.0, ML: 0.2, DV: –0.8), dCA1 (AP: –1.8, ML: 1.4, DV: –1.3), and vCA1 (AP: –3.0, ML: 3.5, DV: –3.3).

Following surgery, each animal was housed individually in a transparent Plexiglas cage with free access to water and food. When these genetic manipulations were combined with electrophysiological recording, a surgery for electrophysiological recording was performed two weeks after the surgery for the genetic manipulations. During the recording and genetic manipulations, the animals did not show any sign of stress or discomfort.

### VNS and electrophysiological recording

To apply VNS to mice, the socket was connected to an electrical stimulator through a wire. Stimulation (VNS; 0.8 mA, 0.1 ms pulse width; 20 Hz; 30 s duration) was performed every 1 min for 3 hours in a day. These individual parameters were set to be in the same ranges as the parameters used in previous studies ^4,6,14,82^.

For c-Fos immunostaining, the mice were transcardially perfused 90 min after VNS.

For LFP recordings, the mice were connected to the recording equipment via Cereplex M (Blackrock Microsystems), a digitally programmable amplifier, which was placed close to the head. The output of the headstage was conducted to the Cereplex Direct recording system, a data acquisition system, via a lightweight multiwire tether and a commutator. All electrophysiological signals were sampled at 2 kHz and low-pass filtered at 500 Hz.

### fMRI

More than one week after the surgery, the mice were first habituated to a custom-made MRI-compatible restraint device without anesthesia by securing the head bar to the device for at least five days. The device was optimized for adjusting the space in the bore of the 7 T magnet with a 23-mm diameter radio frequency volume coil for mice.

fMRI measurements were performed on a Bruker PharmaScan 7T system (Bruker Biospin) with a transmit-receive circularly polarized 23 mm volume coil. For awake-mouse experiments, mice were lightly anesthetized with isoflurane (4–5%) for < 2 min to be placed in the awake-mouse restraint device. BOLD-fMRI was performed more than 30 min after the termination of anesthesia. As reference images for brain structures, T2-weighted images were obtained using the TurboRARE (Rapid Acquisition with Relaxation Enhancement) sequence with the following parameters: repetition time/echo time (TR/TE) = 5500/32.5 ms; field of view (FOV) = 17.5 × 17.5 mm2; image matrix = 256 × 256, number of slices = 48; slice thickness = 0.3 mm; number of averages = 3. For resting-state fMRI, T2*-weighted multislice gradient-echo echo-planar imaging (EPI) was conducted with scan parameters as follows: repetition time/echo time (TR/TE) = 1000/15 ms; field of view (FOV) = 16.4 × 10 mm^2^; image matrix = 82 × 50, number of slices = 20; slice thickness = 0.4 mm; number of averages = 1; bandwidth =250 kHz; flip angle = 30°; number of volumes = 1860 (31 min).

### Fiber Photometry

A fiber-coupled blue LED (470 nm, M470F4, Thorlabs) and yellow LED (M590F3, Thorlabs) excited NE2m and jRGECO1a, respectively, with excitation power set to 25 μW at the fiber tip. The excitation and emission pathways were configured with specific optical components: For 470 nm excitation, light passed through a bandpass filter (86-341, 466/40 nm, Edmund Optics) and was reflected by a dichroic mirror (67-064, 506 nm, Edmund Optics). For 590 nm excitation, light passed through a bandpass filter (84-085, 586/20 nm, Edmund Optics) and was reflected by a dichroic plate (86-321, 605 nm, Edmund Optics). Green fluorescence signals were filtered through a dichroic plate (67-066, 562 nm) and bandpass filter (67-016, 520/36 nm) to isolate 513-538 nm emissions and recorded using a PMT (CR131, Beijing Hamamatsu). Red fluorescence signals were filtered through a bandpass filter (84-087, 628/32 nm) to isolate 612-644 nm emissions and recorded using a PMT (CR316-02, Beijing Hamamatsu).

Signals were digitized using a data acquisition board (COME2-FTR-PYP, Lucir) and recorded using pyPhotometry software at a 100 Hz sampling rate. Data were analyzed using custom Python scripts. Fluorescence traces were first smoothed with a Savitzky–Golay filter (window of 4 s; polynomial order of 20). The filtered traces were then corrected for photobleaching by regression analysis, and ΔF/F was calculated using the mean fluorescence during the first 30 s as the baseline. The root mean square (RMS) of the fluorescence signal was calculated in 1-s windows. jRGECO1a events were defined as fluorescence transients exceeding 1.5 SD above the baseline.

### Chemogenetic manipulation

For chemogenetic experiments to inhibit the NG-cNTS-LC pathway, after at least two weeks following the surgery of guide tube implantation, injection was performed in a home cage. The dummy cannula was removed from the guide tube and replaced with a plastic injection cannula with a diameter of 0.34 mm so that the tip of the injection cannula reached above the LC. The other side of the injection cannula was connected by polyethylene tubing to a 50-μl syringe mounted in an infusion pump (KDS LEGATO101, Muromachi, Japan). DCZ (2.0 μM; #7193, Tocris) dissolved in PBS was infused for 2 min into the LC at a rate of 150 nl/min. Then, the dummy cannula was inserted into the guide tube again.

For chemogenetic experiments to manipulate the VNS-related DMN network, mice received an intraperitoneal injection of 4-hydroxytamoxifen (4-OHT; 100 mg/kg; Sigma-Aldrich, USA; dissolved in a mixture of ethanol and sunflower oil (1:9 v/v) at a final concentration of 10 mg/mL) 1 h before VNS to induce CreER-mediated recombination in c-Fos-expressing neurons. For subsequent chemogenetic manipulation of the trapped neuronal ensembles, mice received either an intraperitoneal injection of DCZ (1 mg/kg; Tocris Bioscience) 30 min before VNS for chemogenetic inhibition or water-soluble DCZ (30 mg/mL) ad libitum in the drinking water for several days prior to behavioral testing for chemogenetic activation. Mice lacking correctly targeted fluorescent protein expression were excluded from subsequent analyses.

### Y-maze spontaneous alternation test

The Y-maze apparatus consisted of three identical black Plexiglas arms (40 × 5 × 15 cm) arranged in a Y-configuration. Mice were placed in the center of the maze and allowed to explore freely for 7 min. Spontaneous alternation behavior was defined as consecutive entries into three different arms. The number of actual alternations was recorded throughout the testing period. Maximum possible alternations were calculated as the total number of arm entries minus 2. The alternation rate was calculated using the following formula: (Actual alternations / Maximum alternations) × 100.

### Novel object recognition test

Mice were habituated on an empty platform (30 × 30 × 30 cm^3^) for at least two days prior to testing. During the first session (learning session), mice were allowed to explore two identical objects on the platform for 10 min. The second session (test session) was conducted 24 hours after the learning session. In the 5-min test phase, mice explored the platform containing one familiar object from the learning session and one novel object. In the test session, the time spent exploring each object was quantified and the discrimination index was calculated according to the following formula:

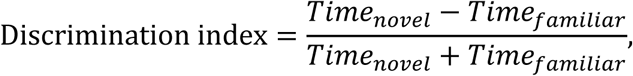

where *Time_novel_* and *Time_familiar_* were the time spent exploring the novel object and the familiar object, respectively. Object exploration was defined as the animal being positioned near the object (distance ≤2 cm) or touching/smelling the object. Simply being placed on the object was not considered exploration.

### Social interaction test

SI tests were performed inside a dark room in a square-shaped box (41.8 cm × 41.8 cm) enclosed by walls 27 cm in height. A wire-mesh cage (5.5 cm × 10 cm × 24 cm) was centered against one wall of the arena during both no-target and target sessions. The interaction zone was defined as a rectangular area (13.5 cm × 24 cm) extending 8 cm around each mesh cage. Corner zones were defined as 9.0 cm × 9.0 cm areas in each corner of the apparatus against the cage. Each SI test included two 150-s sessions (separated by an intersession interval of 30 s) without and with a target C57BL/6N mouse present in the mesh cage, termed no target and target sessions, respectively. In the no target session, a test C57BL/6J mouse was placed in the box and allowed to freely explore the environment. The C57BL/6J mouse was then removed from the box, and a target mouse was next introduced into the mesh cage. The design of the cage allowed the mouse to fit its snout and paws through the mesh cage but not to escape from the cage. In the target session, the same C57BL/6J mouse was placed beside the wall opposite to the mesh cage. A social interaction (SI) ratio was computed as the ratio of time spent in the interaction zone in the target session to the time spent there in the no-target session. To prevent olfactory cues from influencing behavior, the behavioral apparatus was thoroughly cleaned with 70% ethanol between each testing session.

### Six-arm radial maze task

A semiautomated black Plexiglas 8-arm radial maze with a central octagonal platform and eight arms, each 21 cm long and 6.2 cm wide with a wall height of 9 cm, radiating symmetrically outward, was used. Throughout the task, two opposite arms were fully blocked with walls, and behavioral testing was performed using the remaining six arms. Chocolate milk reward (50 μl) was placed at the distal end of the six arms.

In a trial, two randomly selected arms were first blocked with barriers, and the mouse was allowed to freely access the remaining four arms (forced phase). The blocked arms were pseudorandomly arranged with no consecutive occurrences. After the mouse visited four arms and consumed all rewards at four arms, the two barriers were removed, and the mouse was allowed to freely access all six arms (choice phase). There was no delay between the forced phase and choice phase. During the choice phase, the mouse was expected to visit the remaining two arms and obtain the remaining rewards without reentering arms in which the reward had already been consumed. The optimal strategy for solving the maze was to enter each arm only once per trial, and reentries into previously visited arms were considered errors. A trial ended when the mouse had retrieved the reward from all six arms. After an intertrial interval of 2 min, the next trial was initiated. The two blocked arms in the forced phase differed from trial to trial so that the mouse could not adopt a stereotyped strategy to solve the task. The mouse performed 5 trials in a day.

For training of this task, mice underwent three days of food restriction (body weight maintained at >85%), followed by daily 10-min habituation sessions during which each mouse was placed on the central platform with free access to all six arms. During the five-day habituation stage, all mice entered each arm at least once and consumed the chocolate milk reward at the end of each arm within the 10-min period. Task performance was tested beginning the following day.

### Immunohistological analysis

The mice were overdosed with isoflurane, perfused intracardially with 4% paraformaldehyde in PBS and decapitated. After dissection, the brains were fixed overnight in 4% paraformaldehyde and equilibrated with 30% sucrose in PBS for an overnight each. Frozen coronal sections (20–50 μm) were cut using a microtome or cryostat.

For c-Fos immunostaining, the slices were rinsed with 1% bovine serum albumin (BSA) in PBS for 10 min three times, permeabilized with 0.1% Triton X-100 in PBS for 10 min, and then incubated with 1% BSA in PBS for 60 min. The slices were then incubated overnight at 4°C with primary antibodies (1:1000 in 1% BSA/PBS): anti-c-Fos (Encor Biotechnology) and anti-DsRed (632496; Takara Bio). After washing, sections were incubated overnight at 4°C with Alexa 488 anti-mouse IgG (1:500; Invitrogen) in the dark. For EGFP, mCherry, and Cre immunostaining, sections were blocked for 1 hour at room temperature in blocking solution (5% BSA, 0.3% Triton-X in PBS) with gentle agitation. Sections were incubated overnight at 4°C with rabbit anti-GFP (1:1000; 598, MBL), rabbit anti-DsRed (1:500; 632496, Takara Bio), and mouse anti-Cre antibodies (1:1000; MAB3120, Sigma-Aldrich), followed by 90-min room temperature incubation with Alexa 488 anti-rabbit IgG (1:500; Jackson Laboratory), Alexa 594 anti-rabbit IgG (1:500; Jackson Laboratory), and Alexa 594 anti-mouse IgG (1:500; Jackson Laboratory). All sections were washed with PBS, mounted using mounting medium (Vector Laboratories), and imaged using a BZ-X800 fluorescence microscope (Keyence).

c-Fos-positive cells were quantified using the hybrid cell count function. In addition to the brain regions presented in Fig. 1g and 1h, the following regions were also analyzed: temporal association cortex (TeA), nucleus ambiguus (AmbC), median raphe (MR), substantia nigra (SubN), ventral tegmental area (VTA), paraventricular nucleus of the hypothalamus (PVN), bed nucleus of the stria terminalis (BNST), basal forebrain (BF), caudate-putamen complex (CPu), primary and secondary visual cortex (V1 and V2), somatosensory cortex (S1 and S2), motor cortex (M1 and M2), and orbitofrontal cortex (OFC), all of which showed no significant differences (P > 0.05).

After quantifying axon density from LC_DBH,non_ neurons and axon length from LC_DBH,VNS_ neurons, LC_DBH,VNS_ axon fraction in each region was quantified. Assuming that LC neuron projections were entirely distributed among the analyzed brain regions, the fraction of axon density from LC_DBH,non_ neurons in each brain region was calculated by dividing by the total sum across regions (i.e. LC_DBH,non_/∑LC_DBH,non_), representing the degree to which LC_DBH,non_ neurons project preferentially to a given region relative to all regions receiving projections. Similarly, the fraction of LC_DBH,VNS_ axonal length in each brain region was calculated by dividing by the total summed length across regions (i.e. LC_DBH,VNS_/∑ LC_DBH,VNS_). In each brain region, LC_DBH,VNS_ axon fraction was computed as the ratio of these two values (i.e. (LC_DBH,VNS_/∑LC_DBH,VNS_) / (LC_DBH,non_/∑LC_DBH,non_)).

### Spatial transcriptomics using Xenium

For Xenium in situ gene expression analysis, brains were transcardially perfused with 4% paraformaldehyde (PFA), followed by paraffin embedding. Formalin-fixed paraffin-embedded (FFPE) tissues were sectioned at 5-μm thickness and mounted onto Xenium slides according to the Xenium In Situ for FFPE Tissue Preparation Guide (CG000578, Rev F, 10x Genomics). Deparaffinization and decrosslinking were performed according to the Xenium In Situ for FFPE Deparaffinization and Decrosslinking protocol (CG000580, Rev E, 10x Genomics). In situ gene expression analysis was performed using the Xenium Mouse Brain Gene Expression Panel (1000462, 247 genes) together with the Xenium Add-on Custom 1 to 50 Gene Panel (1000652, 50 genes). Probe hybridization, ligation, amplification, autofluorescence quenching, and imaging were performed according to the manufacturer’s instructions. Cell segmentation was performed using the Xenium cell segmentation staining workflow according to the Xenium In Situ Gene Expression with Cell Segmentation Staining User Guide (CG000749, Rev B, 10x Genomics). Slides were processed and imaged using the Xenium Analyzer (10x Genomics) according to the Xenium Analyzer User Guide (CG000584, Rev K). The resulting spatially resolved transcript data and cell segmentation information were used for subsequent analyses. A complete list of the 297 target genes (design ID: 3ZGW8J) included in the pre-designed Xenium Mouse Brain Gene Expression Panel and the Xenium Add-on Custom 1 to 50 Gene Panel is provided in Supplementary Data 1.

### Spatial transcriptomic data analysis

The detection patterns of RNA molecules were visualized using Xenium Explorer (version 1.0, 10x Genomics). Xenium spatial transcriptomic data were analyzed using R with Seurat. Brain regions of interest were manually defined in Xenium Explorer, and the coordinates of cells within each region were extracted for subsequent analyses. For cellular clustering analysis, principal component analysis followed by graph-based clustering was performed. t-SNE was used to visualize cellular distributions and the identified clusters. Glutamatergic, GABAergic, and noradrenergic cell populations were identified based on the expression of *Slc17a7* or *Slc17a6*, *Gad1* or *Gad2*, and *Th*, respectively, as appropriate for each brain region. The spatial distribution of *Fos* transcripts was subsequently examined across the identified clusters.

### In situ hybridization

Probe hybridization and HCR amplification procedures to visualize the localization of DBH-coding mRNA were conducted following the protocol previously described ^83^. The mounting sections were washed with PBS containing 0.2% Triton (PBST) and immersed in 3% H_2_O_2_ in methanol for 10 min, followed by PBST washing for 5 min twice. After washing, the sections were prehybridized for 5 min at 37°C in a hybridization buffer containing 10% dextran sulfate, 0.5× SSC, 0.1% Tween 20, 50 µg/ml heparin, and 1× Denhardt’s solution. The sections were treated with another hybridization solution containing a mixture of 10 nM DBH-probes and incubated overnight at 37°C.

For fluorescent staining, the sections were bleached by a light-emitting diode (LED) illuminator TiYO (Nepa Gene Co., Ltd., Chiba, Japan) for 60 min in PBST to quench autofluorescence after washing.

For HCR amplification, S72 and A161 hairpin DNA pairs were used (Nepa Gene). These hairpin DNA solutions were separately heated to 95 °C for 1 min and then gradually cooled to 65°C for 15 min and to 25°C for 40 min before use. The sections were incubated in an amplification buffer (10% dextran sulfate in 8× SSC, 0.2% Triton X-100, and 100 mM MgCl_2_) for 5 min and then immersed in another amplification buffer containing 60 nM hairpin DNA pairs for 2 h at 25°C. The samples were washed with PBST three times for 5 min.

### Histological analysis for confirmation of electrode placement

To confirm locations of LFP electrodes in the brain, serial sections were mounted and processed for cresyl violet staining. For cresyl violet staining, the slices were rinsed in water, stained with cresyl violet, and coverslipped with Permount. The positions of all electrodes were confirmed by identifying the corresponding electrode tracks in histological tissue.

### fMRI data analysis

Resting-state functional MRI images were preprocessed with AIDAmri, which is a toolbox dedicated to mouse brains using adapted FMRIB Software Library (FSL) tools. Here, datasets underwent bias correction, brain extraction, 3D motion correction, slice-time correction, and registration to a customized brain reference atlas based on the Allen Mouse Brain Atlas. The first six minutes of the time series were removed to account for thermal gradient equilibration. The global signal was then calculated as the average of the percent change time series across all voxels within the brain for each scan. Global signal regression was implemented by regressing out the global signal from each voxel time series to eliminate global noise driven by motion and respiration. A temporal band-pass filter (0.01–0.1 Hz) was then applied. As a result, BOLD signal time series were obtained for specific regions of interest (ROIs) pre-defined within the DMN and SN. The DMN ROIs consisted of the "PL", "IL", "RSC", "PPC", "dCA1", and "vCA1", while the SN ROIs consisted of the "aIC", "pIC", "BLA", and "CeA" (each in both left and right hemispheres). To estimate within-network functional connectivity (FC), Pearson’s correlation coefficients (r) were calculated between the time series of these pre-defined brain regions for each animal. These correlation coefficients were then transformed into z-values using Fisher’s r-to-z transformation to improve normality. The functional connectivity for each network was determined by averaging the z-values of all pairwise correlations between ROIs belonging to that specific network. Furthermore, to evaluate the magnitude of connectivity regardless of the correlation direction (positive or negative), absolute z-values were also calculated and averaged within each network to determine the mean absolute z-values.

The ALFF analysis was carried out using MATLAB. The preprocessed time series of each voxel was transformed into the frequency domain with a Fast Fourier Transform, and the power spectrum was then obtained. We measured ALFF by obtaining the root mean square of the signal across 0.01–0.08 Hz for each voxel. Because ALFF can be susceptible to signal fluctuations caused by physiological noise unrelated to brain activity, we applied a standardization procedure in which the ALFF of each voxel was divided by the global mean ALFF within each animal, resulting in an index called mean ALFF (mALFF). The mALFF value for each brain region was obtained by averaging the voxel-wise mALFF values within each ROI defined by atlas registration.

### LFP analysis

To compute the time-frequency representation of LFP power, brain LFP signals were convolved using complex Morlet wavelet transformation by the MATLAB. The signals that included apparent electrical noise due to physical striking of the animal’s head to the walls were manually removed. When data were obtained from multiple electrodes in a mouse, an electrode was selected so that each mouse contributed one value to the analysis. Regional LFP power was expressed as ratios relative to the corresponding control condition (e.g., post/pre or vagotomy/naive). For inter-regional coherence analysis, coherence between LFP signals recorded from different brain regions was calculated across frequencies. Coherence values were expressed as ratios relative to the corresponding control condition. For hippocampal ripple detection, hippocampal LFP signals were bandpass filtered at 150–250 Hz, and the RMS power was calculated in 10-ms bins. Ripple events were detected when the RMS power exceeded 3 SD above the mean prestress power for at least 15 ms. Periods of massive movement identified from EMG were excluded from the analysis.

### Statistical analysis

All data were presented as the mean ± standard error of the mean (SEM), unless otherwise specified, and were analyzed using MATLAB. For comparisons of c-Fos expression between mouse groups, 10,000 bootstrap samples were generated for each group by resampling with replacement. The mean of each bootstrap sample was calculated, and Student’s *t*-test was applied to the resulting bootstrap datasets. For comparisons between mouse groups, data are displayed as boxplots and analyzed using the Mann–Whitney U test. For comparisons of physiological measures within individual mice, individual data points are displayed together with the sample mean and SEM and analyzed using a paired *t*-test after confirming data normality using the Shapiro–Wilk test. Multiple-group comparisons were performed with Bonferroni post hoc correction. Multiple comparisons were corrected using the Benjamini–Hochberg false discovery rate (FDR) procedure. For the six-arm radial maze task, between-group differences at each testing day were assessed using Welch’s *t*-test with Benjamini–Hochberg FDR correction, and longitudinal changes across testing days were analyzed using repeated-measures ANOVA. The null hypothesis was rejected at the *P* < 0.05 level.

## Data Availability

Source data are provided with this paper.

## Acknowledgements

This work was supported by KAKENHI (24K21735) from the Japan Society for the Promotion of Science (JSPS), grants (JP21zf0127004) from the Japan Agency for Medical Research and Development (AMED), grants from the Japan Science and Technology Agency (JST) (JPMJCR21P1; JPMJMS2292; JPMJKP25Y8) to T. Sasaki; and KAKENHI (24K18233) from JSPS, a grant from JST FOREST Program (JPMJFR231S), a JSPS Research Fellowship for Young Scientists, the Cell Science Research Foundation, the Lotte Foundation, and the Naito Foundation to N. Kuga; and a grant (JP24wm0625116) from AMED to K. Seiriki; and KAKENHI (24K18360) from JSPS and the Konica Minolta Imaging Science Encouragement Award from Konica Minolta Science and Technology Foundation to H. Igarashi; KAKENHI (JP26H02406) from JSPS, a grant (JP25wm0625521) from AMED, and a grant from the Takeda Science Foundation to H. Hashimoto; and the program for Multidisciplinary Frontier Brain and Neuroscience Discoveries (Brain/MINDS 2.0) from AMED (JP23wm0625001) to H. Hirai.

## Author contributions

M.Y., T.T., N.K., and T.S. designed the study. M.Y., T.T., R.S., Z.X., T.K., Y.T., and N.K. performed histological analysis. F.F., T.U., and A.S. performed fMRI measurements. H.K., W.X., T.X., and N.K. acquired and analyzed the electrophysiological data. G.I., S.K., W.H., .M.M., Y.O., and N.A. performed spatial transcriptomic analysis. M.Y. and H.I. performed fiber photometry. Y.I. provided transgenic mouse lines and a technique for virus injection into the nodose ganglion. K.S. and H.Ha. provided virus vectors (AAV9-mDBH-tTA-SV40pA, AAV9-TRE-DIO-Gap43pal-mScarlet-WPRE-hGHpA, and AAV9-TRE-Gap43pal-mScarlet-WPRE-hGHpA). A.K. and H.Hi. provided a virus vector (AAV9-CaMKII-GFP-P2A-Cre). T.T., F.F., T.K., and N.K. acquired the behavioral data. M.Y. and T.S. prepared the figures. N.K. and T.S. wrote the main manuscript text. All the authors reviewed the manuscript text.

---

**Supplementary Figure 1.**
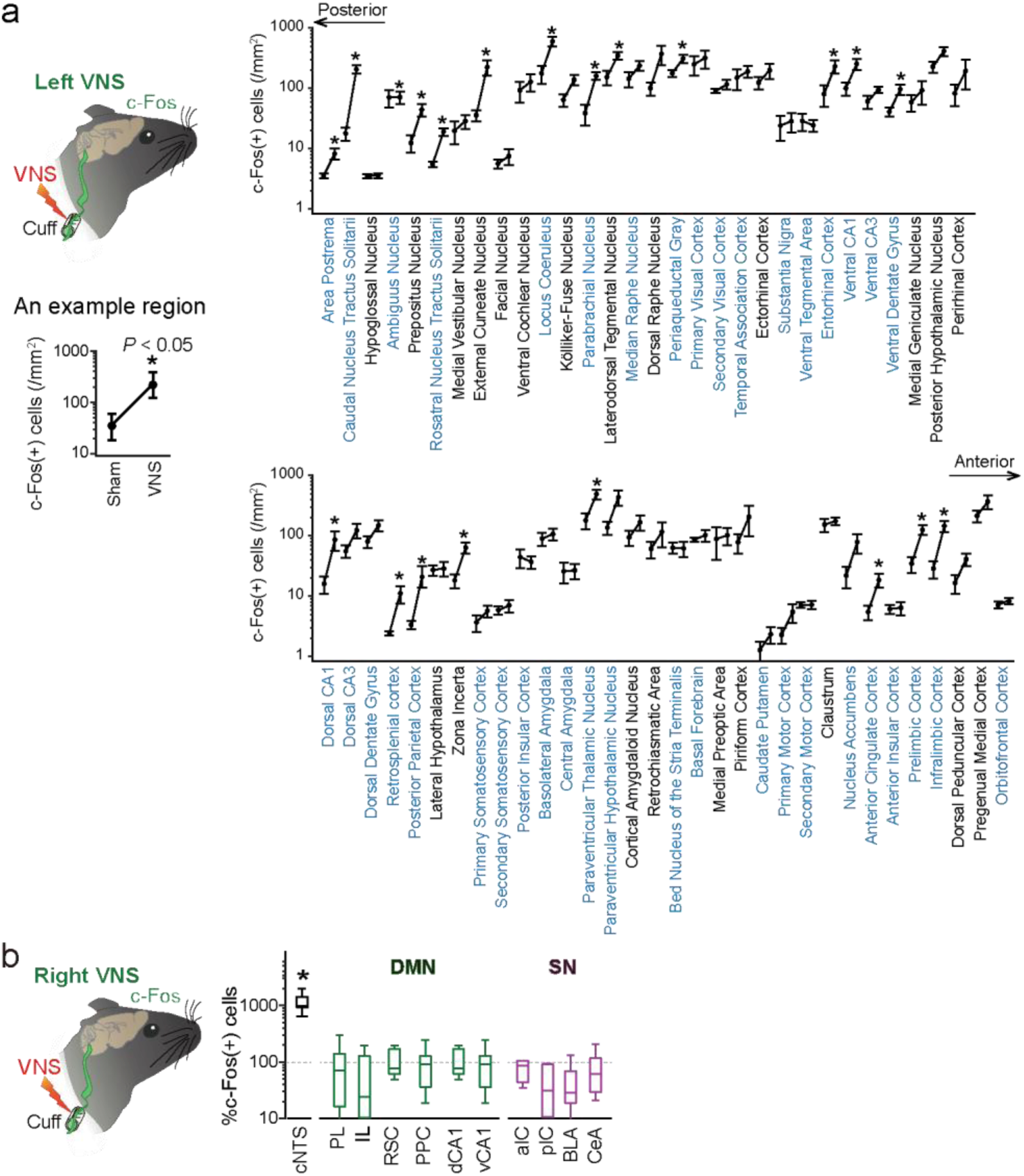
Analysis of brain regions activated by VNS. **a** The numbers of c-Fos-positive neurons were compared between VNS (left plot) and sham (right plot) groups across all 65 brain regions analyzed (*n* = 7 sham and 7–9 VNS mice). Data are presented as mean ± SEM. Brain region names highlighted in Fig. 1g and 1h are colored in blue. * indicates a significant increase (*P* < 0.05), defined from mean values of 10000 bootstrapped samples. **b** The percentages of c-Fos-positive neurons after right VNS in each brain region, compared with sham control (*n* = 7–9 sham and 11 VNS mice). Box plots show the center line as the median, box limits as the 25th percentile and the 75th percentile, and whiskers as minimum to maximum values that are not outliers. \**P* < 0.05, defined from mean values of 10000 bootstrapped samples.

**Supplementary Figure 2.**
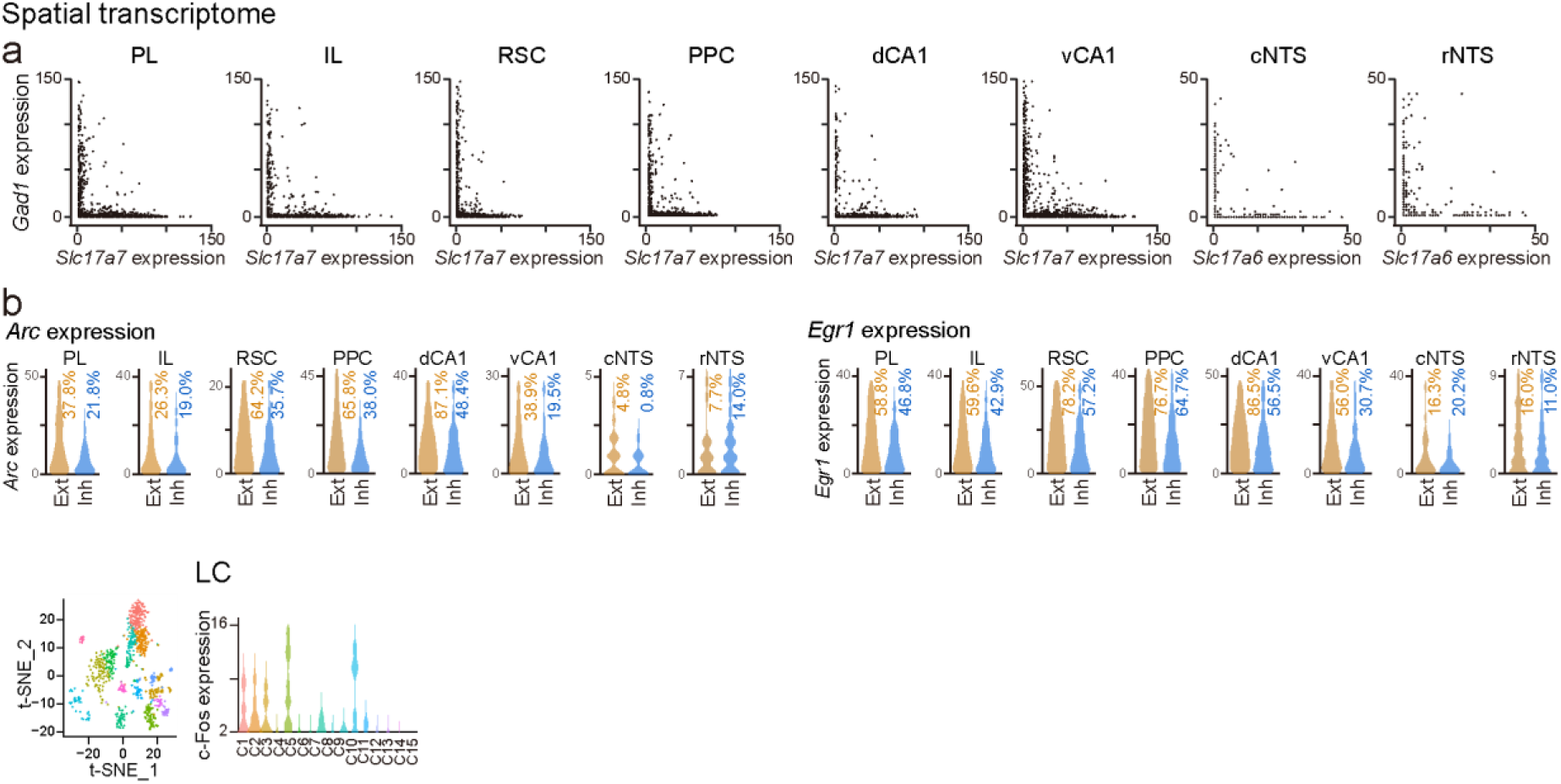
Supplementary results for spatial transcriptomics. **a** Comparison of dot counts between *Slc17a7 or Slc17a6* (Glutamatergic marker) and *Gad1* (GABAergic marker) in each brain region. **b** Distributions of the number of *Arc* or *Egr1* transcript dots detected per cell in excitatory or inhibitory neurons in each region.

**Supplementary Figure 3.**
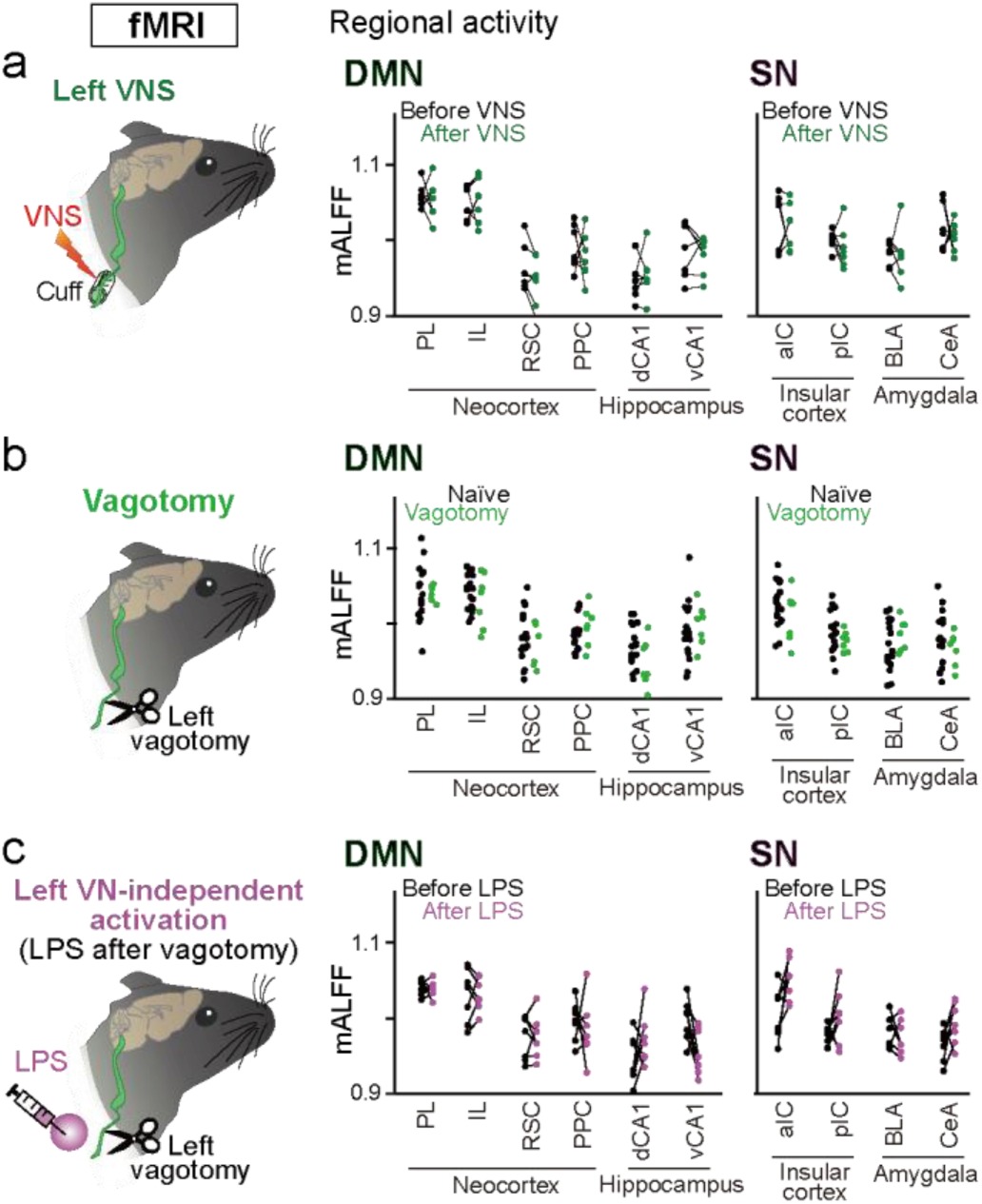
mALFF in individual brain regions in fMRI data. **a** Regional activity as mALFF in individual brain regions before and after VNS (*n* = 7 mice). Each line represents each mouse. **b** Comparisons of mALFF between mice with no vagotomy and vagotomized mice (*n* = 18 and 7 mice). Each dot represents each mouse. **c** Comparisons of mALFF between before and after LPS administration after vagotomy (*n* = 7 mice).

**Supplementary Figure 4.**
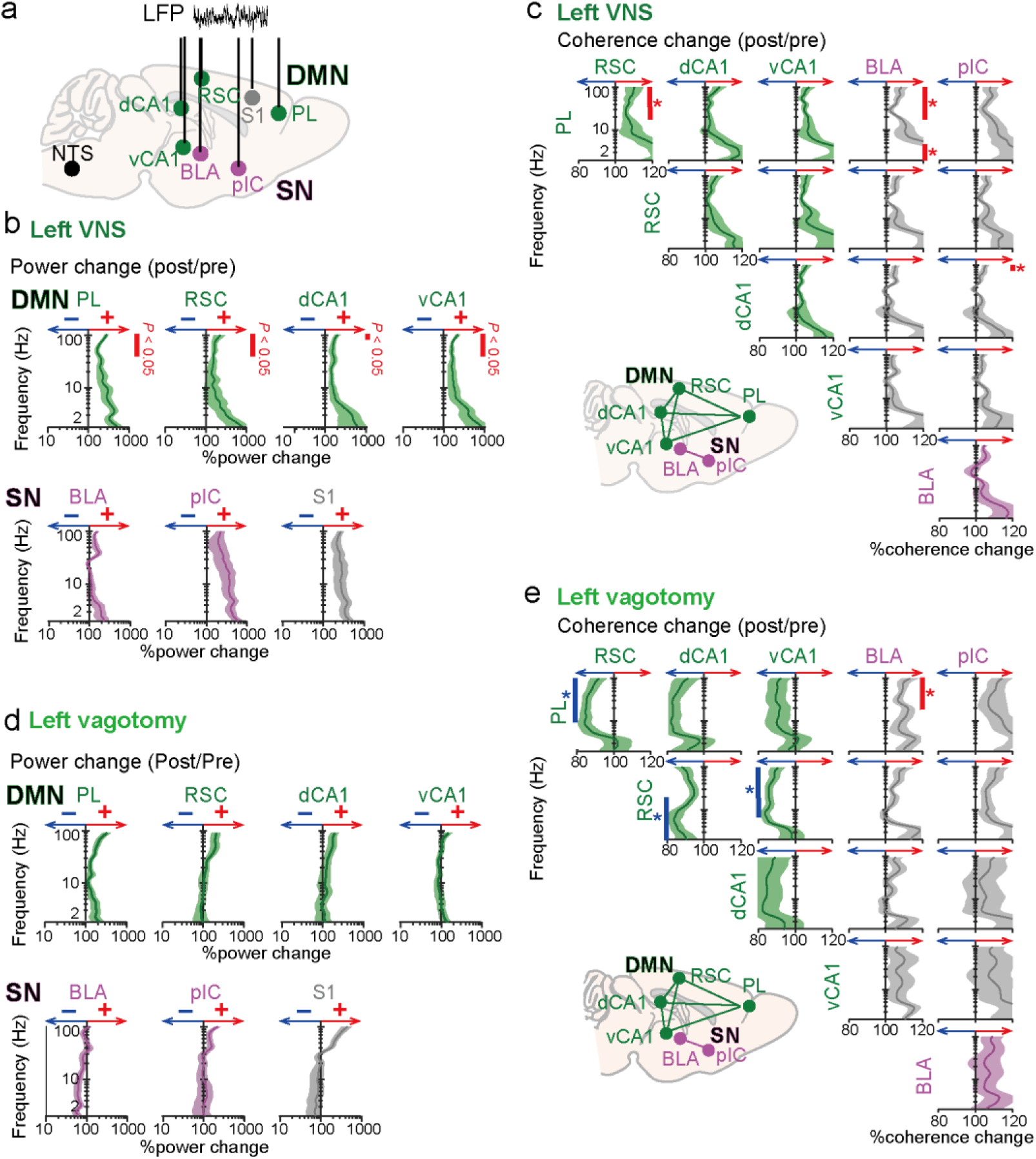
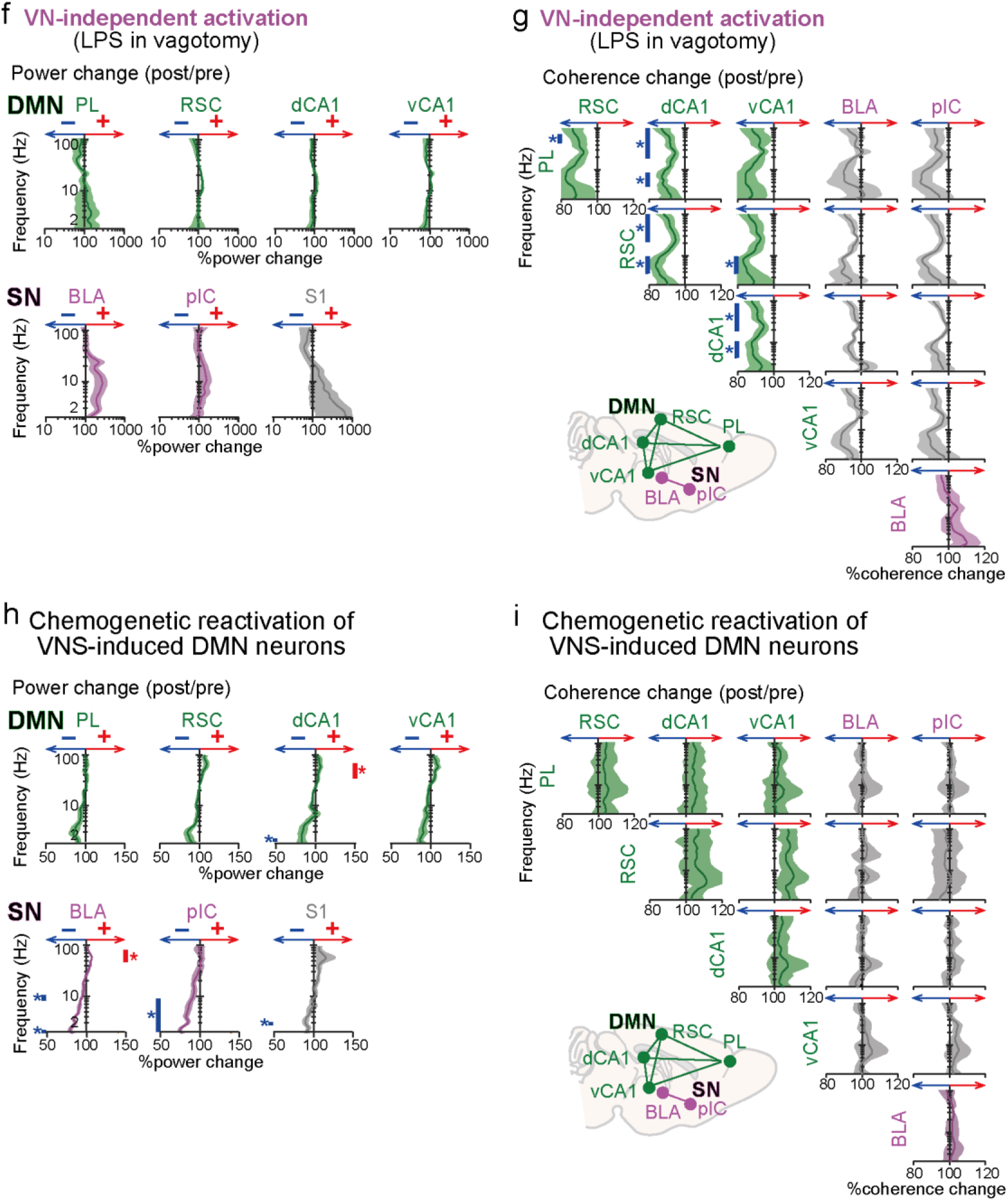
LFP analyses from individual brain regions. **a** LFP recording from the DMN and SN regions. **b** Detailed data for Fig. 2f. Changes in LFP power (2–100 Hz) before (pre) and after (post) VNS in each recorded region (*n* = 9 mice). Shaded areas represent SEM. Red and blue bars indicate frequencies with significant increases and decreases, respectively (*q* < 0.05, paired *t*-test with FDR correction). **c** Detailed data for Fig. 2g. Changes in LFP coherence (2–100 Hz) before (pre) and after (post) VNS between brain region pairs within the DMN and SN. **d** Detailed data for Fig. 2i. Same as b but LFP signals (2–100 Hz) were compared between naïve and vagotomized mice (*n* = 9 and 7 mice). **e** Detailed data for Fig. 2j. Same as c but LFP signals (2–100 Hz) were compared between naïve and vagotomized mice. **f** Detailed data for Fig. 3g. Same as b but LFP signals (2–100 Hz) were compared between before (pre) and after (post) LPS injection after vagotomy (*n* = 7 mice). **g** Detailed data for Fig. 3h. Same as c but LFP signals (2–100 Hz) were compared between before (pre) and after (post) the LPS injection. **h** Detailed data for Fig. 5h. Changes in LFP power (2–100 Hz) before (pre) and after (post) VNS-DMN reactivation in each recorded region (*n* = 6 mice; *q* < 0.05, paired *t*-test with FDR correction). **i** Detailed data for Fig. 5h. Changes in LFP coherence (2–100 Hz) before (pre) and after (post) VNS-DMN reactivation between brain region pairs within the DMN and SN.

**Supplementary Figure 5.**
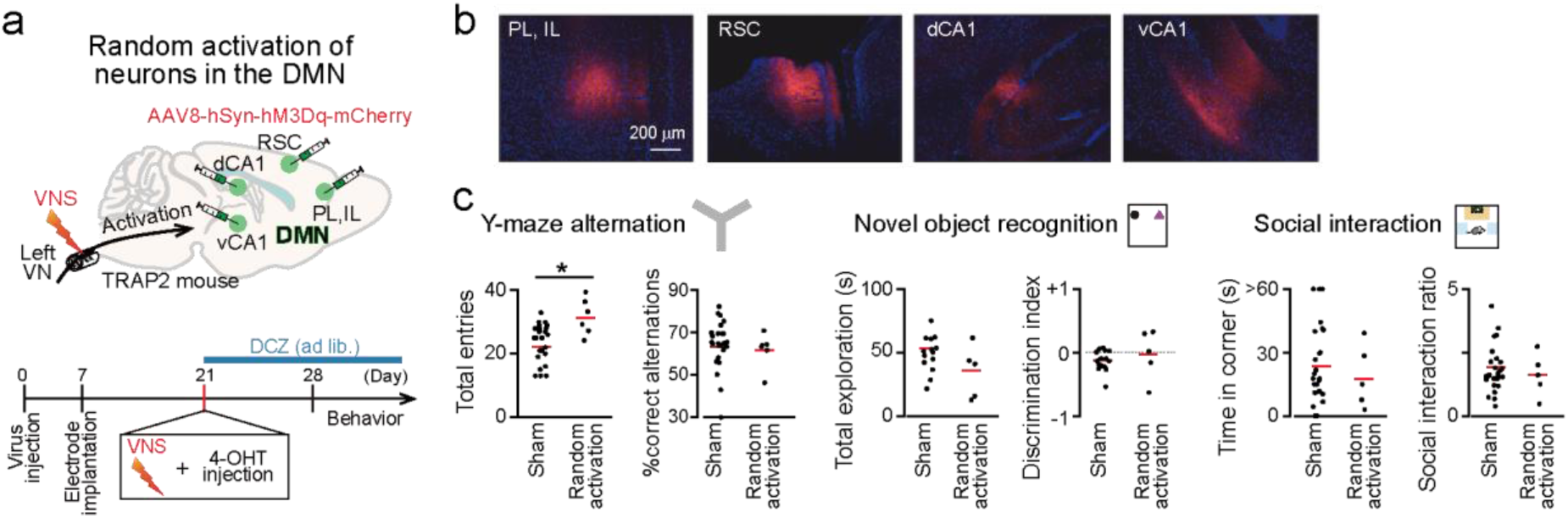
Random activation of neurons in DMN regions does not enhance cognitive behaviors. a,b. The effects of VNS-DMN reactivation may be simply attributable to a general increase in activated neurons within DMN regions. To address this possibility, a random activation group was created by injecting Cre-independent AAV8-hSyn-hM3Dq-mCherry (2.0 × 10¹¹ vg/ml), in which DCZ randomly activated neurons, but without VNS-dependent specificity. The viral titer was adjusted so that the percentage of mCherry-expressing neurons was 10–20%, consistent with that observed from the VNS-DMN reactivation group. **c** (From left to right) Y-maze alternation test performance (*n* = 25 and 6 mice; entry: *Z* = 2.26, *P* = 0.024; alternation: *Z* = 1.03, *P* = 0.30, Mann–Whitney *U* test). Each red line represents the average. Novel object recognition test performance (*n* = 15 and 5 mice; exploration: *Z* = 1.75, *P* = 0.081; index: *Z* = 0.79, *P* = 0.43, Mann–Whitney *U* test). Social interaction test performance (*n* = 27 and 5 mice; corner: *Z* = 0.42, *P* = 0.68; ratio: *Z* = 0.18, *P* = 0.86, Mann–Whitney *U* test).

## References

1 Bravo, J. A. et al. Ingestion of Lactobacillus strain regulates emotional behavior and central GABA receptor expression in a mouse via the vagus nerve. Proc Natl Acad Sci U S A 108, 16050–16055, doi:10.1073/pnas.1102999108 (2011).

2 Liu, Y. & Forsythe, P. Vagotomy and insights into the microbiota-gut-brain axis. Neurosci Res 168, 20–27, doi:10.1016/j.neures.2021.04.001 (2021).

3 Kuga, N. et al. Sniffing behaviour-related changes in cardiac and cortical activity in rats. J Physiol 597, 5295–5306, doi:10.1113/JP278500 (2019).

4 Okonogi, T. et al. Stress-induced vagal activity influences anxiety-relevant prefrontal and amygdala neuronal oscillations in male mice. Nat Commun 15, 183, doi:10.1038/s41467-023-44205-y (2024).

5 Klarer, M. et al. Gut vagal afferents differentially modulate innate anxiety and learned fear. J Neurosci 34, 7067–7076, doi:10.1523/JNEUROSCI.0252-14.2014 (2014).

6 Furmaga, H., Shah, A. & Frazer, A. Serotonergic and noradrenergic pathways are required for the anxiolytic-like and antidepressant-like behavioral effects of repeated vagal nerve stimulation in rats. Biol Psychiatry 70, 937–945, doi:10.1016/j.biopsych.2011.07.020 (2011).

7 Rush, A. J. et al. Vagus nerve stimulation (VNS) for treatment-resistant depressions: a multicenter study. Biol Psychiatry 47, 276–286, doi:10.1016/s0006-3223(99)00304-2 (2000).

8 Austelle, C. W., Cox, S. S., Wills, K. E. & Badran, B. W. Vagus nerve stimulation (VNS): recent advances and future directions. Clin Auton Res 34, 529–547, doi:10.1007/s10286-024-01065-w (2024).

9 Noble, L. J. et al. Effects of vagus nerve stimulation on extinction of conditioned fear and post-traumatic stress disorder symptoms in rats. Transl Psychiatry 7, e1217, doi:10.1038/tp.2017.191 (2017).

10 Neuser, M. P. et al. Vagus nerve stimulation boosts the drive to work for rewards. Nat Commun 11, 3555, doi:10.1038/s41467-020-17344-9 (2020).

11 Lucchi, F., Lloyd, B. & Nieuwenhuis, S. Non-invasive vagus nerve stimulation and the motivation to work for rewards: A replication of Neuser et al. (2020, Nature Communications). Psychophysiology 61, e14484, doi:10.1111/psyp.14484 (2024).

12 Martin, K. A. et al. Vagus nerve stimulation recruits the central cholinergic system to enhance perceptual learning. Nat Neurosci, doi:10.1038/s41593-024-01767-4 (2024).

13 Teckentrup, V. & Kroemer, N. B. Mechanisms for survival: vagal control of goal-directed behavior. Trends Cogn Sci 28, 237–251, doi:10.1016/j.tics.2023.11.001 (2024).

14 Noble, L. J. et al. Vagus nerve stimulation promotes generalization of conditioned fear extinction and reduces anxiety in rats. Brain Stimul 12, 9–18, doi:10.1016/j.brs.2018.09.013 (2019).

15 Clark, K. B. et al. Posttraining electrical stimulation of vagal afferents with concomitant vagal efferent inactivation enhances memory storage processes in the rat. Neurobiol Learn Mem 70, 364–373, doi:10.1006/nlme.1998.3863 (1998).

16 Pena, D. F., Engineer, N. D. & McIntyre, C. K. Rapid remission of conditioned fear expression with extinction training paired with vagus nerve stimulation. Biol Psychiatry 73, 1071–1077, doi:10.1016/j.biopsych.2012.10.021 (2013).

17 Bowles, S. et al. Vagus nerve stimulation drives selective circuit modulation through cholinergic reinforcement. Neuron 110, 2867–2885 e2867, doi:10.1016/j.neuron.2022.06.017 (2022).

18 Suarez, A. N. et al. Gut vagal sensory signaling regulates hippocampus function through multi-order pathways. Nat Commun 9, 2181, doi:10.1038/s41467-018-04639-1 (2018).

19 Thakkar, V. J., Engelhart, A. S., Khodaparast, N., Abadzi, H. & Centanni, T. M. Transcutaneous auricular vagus nerve stimulation enhances learning of novel letter-sound relationships in adults. Brain Stimul 13, 1813–1820, doi:10.1016/j.brs.2020.10.012 (2020).

20 Olsen, L. K., Solis, E., Jr., McIntire, L. K. & Hatcher-Solis, C. N. Vagus nerve stimulation: mechanisms and factors involved in memory enhancement. Front Hum Neurosci 17, 1152064, doi:10.3389/fnhum.2023.1152064 (2023).

21 Szeska, C., Ventura-Bort, C., Giraudier, M. & Weymar, M. A vagal route to memory: evidence from invasive and non-invasive electrical vagus nerve stimulation studies and areas for future clinical application. Front Hum Neurosci 19, 1595737, doi:10.3389/fnhum.2025.1595737 (2025).

22 Collins, L., Boddington, L., Steffan, P. J. & McCormick, D. Vagus nerve stimulation induces widespread cortical and behavioral activation. Curr Biol 31, 2088–2098 e2083, doi:10.1016/j.cub.2021.02.049 (2021).

23 Tierno Lauer, L., et al. The vagus nerve promotes memory in rats via nutrient-induced septo-hippocampal acetylcholine signaling. Nat Commun, doi:10.1038/s41467-026-73896-2 (2026).

24 Smallwood, J. et al. The default mode network in cognition: a topographical perspective. Nat Rev Neurosci 22, 503–513, doi:10.1038/s41583-021-00474-4 (2021).

25 Menon, V. 20 years of the default mode network: A review and synthesis. Neuron 111, 2469–2487, doi:10.1016/j.neuron.2023.04.023 (2023).

26 Raichle, M. E. et al. A default mode of brain function. Proc Natl Acad Sci U S A 98, 676– 682, doi:10.1073/pnas.98.2.676 (2001).

27 Greicius, M. D., Krasnow, B., Reiss, A. L. & Menon, V. Functional connectivity in the resting brain: a network analysis of the default mode hypothesis. Proc Natl Acad Sci U S A 100, 253–258, doi:10.1073/pnas.0135058100 (2003).

28 Uddin, L. Q., Yeo, B. T. T. & Spreng, R. N. Towards a Universal Taxonomy of Macro-scale Functional Human Brain Networks. Brain Topogr 32, 926–942, doi:10.1007/s10548-019-00744-6 (2019).

29 Buckner, R. L., Andrews-Hanna, J. R. & Schacter, D. L. The brain’s default network: anatomy, function, and relevance to disease. Ann N Y Acad Sci 1124, 1–38, doi:10.1196/annals.1440.011 (2008).

30 Alves, P. N. et al. An improved neuroanatomical model of the default-mode network reconciles previous neuroimaging and neuropathological findings. Commun Biol 2, 370, doi:10.1038/s42003-019-0611-3 (2019).

31 Raichle, M. E. The brain’s default mode network. Annu Rev Neurosci 38, 433–447, doi:10.1146/annurev-neuro-071013-014030 (2015).

32 Bartoli, E. et al. Default mode network electrophysiological dynamics and causal role in creative thinking. Brain 147, 3409–3425, doi:10.1093/brain/awae199 (2024).

33 Fang, J. et al. Transcutaneous Vagus Nerve Stimulation Modulates Default Mode Network in Major Depressive Disorder. Biol Psychiatry 79, 266–273, doi:10.1016/j.biopsych.2015.03.025 (2016).

34 Mandino, F. et al. A triple-network organization for the mouse brain. Mol Psychiatry 27, 865–872, doi:10.1038/s41380-021-01298-5 (2022).

35 Whitesell, J. D. et al. Regional, Layer, and Cell-Type-Specific Connectivity of the Mouse Default Mode Network. Neuron 109, 545–559 e548, doi:10.1016/j.neuron.2020.11.011 (2021).

36 Stafford, J. M. et al. Large-scale topology and the default mode network in the mouse connectome. Proc Natl Acad Sci U S A 111, 18745–18750, doi:10.1073/pnas.1404346111 (2014).

37 Chao, T. H. et al. Neuronal dynamics of the default mode network and anterior insular cortex: Intrinsic properties and modulation by salient stimuli. Sci Adv 9, eade5732, doi:10.1126/sciadv.ade5732 (2023).

38 Menon, V. et al. Optogenetic stimulation of anterior insular cortex neurons in male rats reveals causal mechanisms underlying suppression of the default mode network by the salience network. Nat Commun 14, 866, doi:10.1038/s41467-023-36616-8 (2023).

39 Lu, H. et al. Rat brains also have a default mode network. Proc Natl Acad Sci U S A 109, 3979–3984, doi:10.1073/pnas.1200506109 (2012).

40 Nghiem, T. E. et al. Space wandering in the rodent default mode network. Proc Natl Acad Sci U S A 121, e2315167121, doi:10.1073/pnas.2315167121 (2024).

41 Gutierrez-Barragan, D. et al. Unique spatiotemporal fMRI dynamics in the awake mouse brain. Curr Biol 32, 631–644 e636, doi:10.1016/j.cub.2021.12.015 (2022).

42 Mughrabi, I. T. et al. Development and characterization of a chronic implant mouse model for vagus nerve stimulation. Elife 10, doi:10.7554/eLife.61270 (2021).

43 Mendez-Hernandez, R., Braga, I., Bali, A., Yang, M. & de Lartigue, G. Vagal Sensory Gut-Brain Pathways That Control Eating-Satiety and Beyond. Compr Physiol 15, e70010, doi:10.1002/cph4.70010 (2025).

44 Iba, K., Kitano, R. & Iwasaki, Y. Gut sensing of food ingredients and interoception-mediated regulation of feeding and glucose metabolism. Physiol Rep 13, e70636, doi:10.14814/phy2.70636 (2025).

45 Masuda, Y., Ohbayashi, K. & Iwasaki, Y. Comparing the anorexigenic effects and mechanisms of gut-derived GLP-1 and its receptor agonists: insights into incretin-based therapies for obesity. Diabetol Int 16, 448–456, doi:10.1007/s13340-025-00819-9 (2025).

46 Logothetis, N. K., Pauls, J., Augath, M., Trinath, T. & Oeltermann, A. Neurophysiological investigation of the basis of the fMRI signal. Nature 412, 150–157, doi:10.1038/35084005 (2001).

47 Zou, Q. H. et al. An improved approach to detection of amplitude of low-frequency fluctuation (ALFF) for resting-state fMRI: fractional ALFF. J Neurosci Methods 172, 137–141, doi:10.1016/j.jneumeth.2008.04.012 (2008).

48 Buzsaki, G. Hippocampal sharp wave-ripple: A cognitive biomarker for episodic memory and planning. Hippocampus 25, 1073–1188, doi:10.1002/hipo.22488 (2015).

49 Norman, Y., Raccah, O., Liu, S., Parvizi, J. & Malach, R. Hippocampal ripples and their coordinated dialogue with the default mode network during recent and remote recollection. Neuron 109, 2767–2780 e2765, doi:10.1016/j.neuron.2021.06.020 (2021).

50 Kaplan, R. et al. Hippocampal Sharp-Wave Ripples Influence Selective Activation of the Default Mode Network. Curr Biol 26, 686–691, doi:10.1016/j.cub.2016.01.017 (2016).

51 Zingg, B. et al. AAV-Mediated Anterograde Transsynaptic Tagging: Mapping Corticocollicular Input-Defined Neural Pathways for Defense Behaviors. Neuron 93, 33– 47, doi:10.1016/j.neuron.2016.11.045 (2017).

52 Hulsey, D. R. et al. Parametric characterization of neural activity in the locus coeruleus in response to vagus nerve stimulation. Exp Neurol 289, 21–30, doi:10.1016/j.expneurol.2016.12.005 (2017).

53 Yakunina, N., Kim, S. S. & Nam, E. C. Optimization of Transcutaneous Vagus Nerve Stimulation Using Functional MRI. Neuromodulation 20, 290–300, doi:10.1111/ner.12541 (2017).

54 Groves, D. A., Bowman, E. M. & Brown, V. J. Recordings from the rat locus coeruleus during acute vagal nerve stimulation in the anaesthetised rat. Neurosci Lett 379, 174– 179, doi:10.1016/j.neulet.2004.12.055 (2005).

55 Itoi, K. et al. Targeting of locus ceruleus noradrenergic neurons expressing human interleukin-2 receptor alpha-subunit in transgenic mice by a recombinant immunotoxin anti-Tac(Fv)-PE38: a study for exploring noradrenergic influence upon anxiety-like and depression-like behaviors. J Neurosci 31, 6132–6139, doi:10.1523/JNEUROSCI.5188-10.2011 (2011).

56 Manta, S., El Mansari, M., Debonnel, G. & Blier, P. Electrophysiological and neurochemical effects of long-term vagus nerve stimulation on the rat monoaminergic systems. Int J Neuropsychopharmacol 16, 459–470, doi:10.1017/S1461145712000387 (2013).

57 Seiriki, K. et al. Noradrenergic efferent subsystems that gate traumatic social learning. bioRxiv, 2025.2003.2007.641347, doi:10.1101/2025.03.07.641347 (2025).

58 Schwarz, L. A. & Luo, L. Organization of the locus coeruleus-norepinephrine system. Curr Biol 25, R1051–R1056, doi:10.1016/j.cub.2015.09.039 (2015).

59 Uematsu, A. et al. Modular organization of the brainstem noradrenaline system coordinates opposing learning states. Nat Neurosci 20, 1602–1611, doi:10.1038/nn.4642 (2017).

60 Poe, G. R. et al. Locus coeruleus: a new look at the blue spot. Nat Rev Neurosci 21, 644– 659, doi:10.1038/s41583-020-0360-9 (2020).

61 Kebschull, J. M. et al. High-Throughput Mapping of Single-Neuron Projections by Sequencing of Barcoded RNA. Neuron 91, 975–987, doi:10.1016/j.neuron.2016.07.036 (2016).

62 DeNardo, L. A. et al. Temporal evolution of cortical ensembles promoting remote memory retrieval. Nat Neurosci 22, 460–469, doi:10.1038/s41593-018-0318-7 (2019).

63 Guenthner, C. J., Miyamichi, K., Yang, H. H., Heller, H. C. & Luo, L. Permanent genetic access to transiently active neurons via TRAP: targeted recombination in active populations. Neuron 78, 773–784, doi:10.1016/j.neuron.2013.03.025 (2013).

64 Feng, J. et al. A Genetically Encoded Fluorescent Sensor for Rapid and Specific In Vivo Detection of Norepinephrine. Neuron 102, 745–761 e748, doi:10.1016/j.neuron.2019.02.037 (2019).

65 Feng, J. et al. Monitoring norepinephrine release in vivo using next-generation GRAB(NE) sensors. Neuron 112, 1930–1942 e1936, doi:10.1016/j.neuron.2024.03.001 (2024).

66 Zhang, H. et al. Cognitive function and brain activation before and after transcutaneous cervical vagus nerve stimulation in healthy adults: A concurrent tcVNS-fMRI study. Front Psychol 13, 1003411, doi:10.3389/fpsyg.2022.1003411 (2022).

67 Kraus, T. et al. BOLD fMRI deactivation of limbic and temporal brain structures and mood enhancing effect by transcutaneous vagus nerve stimulation. J Neural Transm (Vienna) 114, 1485–1493, doi:10.1007/s00702-007-0755-z (2007).

68 Dietrich, S. et al. A novel transcutaneous vagus nerve stimulation leads to brainstem and cerebral activations measured by functional MRI. Biomed Tech (Berl) 53, 104–111, doi:10.1515/BMT.2008.022 (2008).

69 Monti, D. A. et al. Changes in brain functional connectivity associated with transcutaneous auricular vagus nerve stimulation in healthy controls. Front Hum Neurosci 19, 1531123, doi:10.3389/fnhum.2025.1531123 (2025).

70 Joo, H. R. & Frank, L. M. The hippocampal sharp wave-ripple in memory retrieval for immediate use and consolidation. Nat Rev Neurosci 19, 744–757, doi:10.1038/s41583-018-0077-1 (2018).

71 Zhao, Q. et al. A multidimensional coding architecture of the vagal interoceptive system. Nature 603, 878–884, doi:10.1038/s41586-022-04515-5 (2022).

72 Jayaprakash, N. et al. Organ- and function-specific anatomical organization of vagal fibers supports fascicular vagus nerve stimulation. Brain Stimul 16, 484–506, doi:10.1016/j.brs.2023.02.003 (2023).

73 Han, W. et al. A Neural Circuit for Gut-Induced Reward. Cell 175, 665–678 e623, doi:10.1016/j.cell.2018.08.049 (2018).

74 Shaffer, C., Barrett, L. F. & Quigley, K. S. Signal processing in the vagus nerve: Hypotheses based on new genetic and anatomical evidence. Biol Psychol 182, 108626, doi:10.1016/j.biopsycho.2023.108626 (2023).

75 Schappe, M. S. et al. A vagal reflex evoked by airway closure. Nature 627, 830–838, doi:10.1038/s41586-024-07144-2 (2024).

76 Iwasaki, Y. & Yada, T. Vagal afferents sense meal-associated gastrointestinal and pancreatic hormones: mechanism and physiological role. Neuropeptides 46, 291–297, doi:10.1016/j.npep.2012.08.009 (2012).

77 Conio, B. et al. Opposite effects of dopamine and serotonin on resting-state networks: review and implications for psychiatric disorders. Mol Psychiatry 25, 82–93, doi:10.1038/s41380-019-0406-4 (2020).

78 Aguilar, D. D. & McNally, J. M. Subcortical control of the default mode network: Role of the basal forebrain and implications for neuropsychiatric disorders. Brain Res Bull 185, 129–139, doi:10.1016/j.brainresbull.2022.05.005 (2022).

79 Cunningham, J. T., Mifflin, S. W., Gould, G. G. & Frazer, A. Induction of c-Fos and DeltaFosB immunoreactivity in rat brain by Vagal nerve stimulation. Neuropsychopharmacology 33, 1884–1895, doi:10.1038/sj.npp.1301570 (2008).

80 Wise, T. et al. Instability of default mode network connectivity in major depression: a two-sample confirmation study. Transl Psychiatry 7, e1105, doi:10.1038/tp.2017.40 (2017).

81 Kuga, N., Abe, R., Takano, K., Ikegaya, Y. & Sasaki, T. Prefrontal-amygdalar oscillations related to social behavior in mice. Elife 11, doi:10.7554/eLife.78428 (2022).

82 Choudhary, R. C. et al. Threshold adjusted vagus nerve stimulation after asphyxial cardiac arrest results in neuroprotection and improved survival. Bioelectron Med 8, 10, doi:10.1186/s42234-022-00092-0 (2022).

83 Tsuneoka, Y. & Funato, H. Modified in situ Hybridization Chain Reaction Using Short Hairpin DNAs. Front Mol Neurosci 13, 75, doi:10.3389/fnmol.2020.00075 (2020).

